# Device encapsulated MSCs for adaptive secretome therapy to effectively target ischaemic heart injury

**DOI:** 10.1101/2025.04.01.646502

**Authors:** Andrew R. Kompa, David W. Greening, Jarmon G. Lees, Anne M. Kong, Jonathon Cross, Ashley Nowland, Ren J. Phang, Saba Naghipour, Yali Deng, Jack R.T. Darby, Lina Mariana, Cameron Kos, Tanya Hall, Andrew Newcomb, James J.H. Chong, Rebecca H Ritchie, Janna L. Morrison, Klearchos K. Papas, Kilian Kelly, Derek J. Hausenloy, Thomas Loudovaris, Shiang Y. Lim

## Abstract

Effective long-term strategies to protect the ischaemic heart remain a significant challenge. Mesenchymal stem cells (MSCs) offer therapeutic potential primarily through their secretome, a bioactive factor-rich milieu with broad beneficial effects. However, existing delivery methods have not been shown to provide sustained benefits. Herein, we introduce an innovative approach for sustained MSC-secretome delivery for long-term cardioprotection. In a rat model of chronic myocardial infarction, Cymerus MSCs, derived from human induced pluripotent stem cells (iPSCs), were encapsulated in a Procyon immunoisolation device and implanted subcutaneously. A human-iPSC-derived engineered cardiac microtissue model was used to simulate ischaemia-reperfusion injury and assess cardioprotective effects in a human context. The MSC-loaded Procyon device significantly improved cardiac function and reduced adverse left ventricular remodelling over a 12-week period. These positive effects were consistent across both young and middle-aged, male and female rats, indicating broad applicability. The encapsulated MSCs remained viable and continuously released therapeutic secretome for 12 weeks. *In vitro*, the MSC secretome protected human engineered cardiac microtissues from simulated ischaemia-reperfusion injury, restoring contractile function, improving cell viability, and reducing oxidative stress. Proteomic analysis of MSCs revealed 179 unique cellular proteins post-implantation, linked to adaptive immune and inflammatory responses as well as wound healing. MSC secretome profiling revealed increased protein diversity associated with tissue repair and immune regulation, suggesting MSCs undergo an adaptive response to ischaemic conditions, enhancing their therapeutic potential. This translational study highlights a clinically viable, minimally invasive method for sustained cardioprotection, harnessing the MSC secretome to address a pivotal gap in current treatments for ischaemic heart disease.

## Introduction

Ischaemic heart disease remains the leading cause of death globally, with acute myocardial infarction (AMI) as a key clinical manifestation ^1^. A major complication following AMI is adverse left ventricular remodelling, which often progresses to heart failure. Although several therapies are available to protect and repair the heart post-AMI, their effectiveness diminishes as the disease advances and the extent of damage increases ^2^. Consequently, there is a critical unmet need for more effective long-term cardioprotective interventions. Stem cell therapy presents a promising opportunity to address this critical unmet need.

Mesenchymal stem cells (MSCs), including those derived from human induced pluripotent stem cells (iPSCs), have shown promising potential to enhance cardiac function and structure, primarily through the paracrine activity of their secretome ^3–5^. This secretome significantly improves the myocardial microenvironment by activating endogenous repair mechanisms; these include enhancing cell survival, promoting angiogenesis, resolving inflammation, and mitigating adverse remodelling ^6,7^. These promising preclinical findings on the stem cell secretome are further reinforced by the ongoing first-in-human Phase 1 clinical trial, SECRET-HF, which specifically evaluates the cardioprotective effect of an extracellular vesicle-enriched secretome derived from cardiovascular progenitor cells generated from human iPSCs. While the trial is still underway, a case study of a patient with non-ischaemic dilated cardiomyopathy who received three intravenous infusions of secretome at three-week intervals showed promising cardiac improvements with no signs of immunogenicity ^8^.

Despite these advancements, clinical studies using MSCs from various sources have only demonstrated modest improvements in cardiac function. The limited success of stem cell therapies is partly due to suboptimal cell delivery methods, which fail to fully harness the therapeutic potential of stem cells for sustained benefits and long-term cardiac repair ^9,10^. Current techniques, whether through intravenous, intracoronary, or direct intramyocardial delivery, often suffer from poor cell retention and high cell death rates in the harsh ischaemic environment, limiting the beneficial effects of the paracrine factors secreted by transplanted cells. These effects are typically short-lived, and the invasive nature of repeat administration poses a significant clinical challenge. While innovative approaches, such as cell encapsulating hydrogels, have shown promise in improving cell retention, they are still invasive and provide only temporary benefits due to their rapid biodegradation and death of the encapsulated MSCs within ^9,10^.

In response to these challenges, we have pioneered an innovative and minimally invasive approach to harness MSCs as a living biodrug for the sustained release of their secretome, offering an effective strategy for long-term cardioprotection. Our method utilised a retrievable immunoisolation device implanted subcutaneously, ensuring both safety and controlled, long-term delivery of MSC-derived therapeutic secretome ^11^. This device features a dual-membrane system: the inner membrane shields encapsulated cells from the host immune system while allowing the diffusion of nutrients and therapeutic outputs (including the secretome), and the outer membrane promotes blood vessel formation. In a preclinical study, subcutaneous implantation of these devices encapsulating human cardiac stem cells improved cardiac function and prevented maladaptive remodelling four weeks after non-reperfused MI in immunocompromised rats ^11^.

Despite these encouraging results, a significant gap remains in understanding the long-term efficacy of stem cell secretome therapies in immunocompetent models of chronic myocardial infarction. Addressing this limitation is essential for clinical translation, as immunocompromised models do not accurately reflect the immune responses seen in human patients, and short-term studies fail to capture the long-term remodelling that follows AMI. To bridge this translational gap, the present study evaluates the long-term effectiveness of MSC secretome-based therapies using clinical-grade MSCs derived from iPSCs in an immunocompetent rat model of chronic reperfused myocardial infarction, as well as in a human iPSC-derived cardiac microtissue model of ischaemia-reperfusion injury. The results show that sustained MSC secretome therapy provides long-term cardioprotection in age-appropriate models of reperfused myocardial infarction in both sexes, as well as in a preclinical human cardiac microtissues. Proteomic analysis further reveals that encapsulated MSCs exhibit an adaptive response by secreting a different combination of paracrine factors to target and modulate cardiac pathologies, thereby enhancing cardioprotection.

## Methods

### Cell culture

#### Cymerus^TM^ MSCs

Cymerus MSCs were generated from clinical-grade human induced pluripotent stem cells (iPSCs) as previously described ^12^ and supplied by Cynata Therapeutics Limited (Victoria, Australia). Upon receipt, the cryopreserved Cymerus MSCs (passage 2) were thawed, seeded on tissue culture flasks pre-coated with human fibronectin (1 mg/mL; Thermo Fisher Scientific, MA, USA) and human collagen I (3 µg/mL; Sigma-Aldrich, MO, USA), and expanded in serum-free expansion medium according to the manufacturer’s instructions. Cymerus MSCs at passages 6-7 were used in the present study.

#### Human iPSC culture and differentiation

The human iPS-Foreskin-2 cell line, kindly provided by James A. Thomson (University of Wisconsin) ^13^ was maintained on vitronectin-coated plates in TeSR-E8 medium (STEMCELL Technologies, Vancouver, Canada) according to the manufacturer’s protocol.

### Engineered cardiac microtissue

To construct the multicellular cardiac microtissue, iPSC-derived cardiomyocytes (3.5×10^4^ cells), endothelial cells (1.25×10^4^ cells), smooth muscle cells (1.25 ×10^3^ cells) and cardiac fibroblasts (1.25×10^3^ cells) were seeded per well into ultra-low attachment round-bottom 96-well plates in spheroid induction medium (cardiomyocyte replating medium, EGM2-MV, SmGM-2, and FGM-3 at a 1:1:1:1 ratio and supplemented with 50 ng/mL VEGF-165 and 10 μM Y-27632), and spun at 150 *g* for 3 minutes. After 48 hours, medium was changed to cardiac spheroid maintaining medium (cardiomyocyte medium, EGM2-MV, SmGM-2, and FGM-3 at a 1:1:1:1 ratio and supplemented with 50 ng/mL VEGF-165). After a further 24 hours, compacted spheroids were embedded in 10 µL growth factor reduced Matrigel and transferred to tissue culture plates coated with anti-adherence rinsing solution (STEMCELL Technologies) containing cardiac spheroid maintaining medium. Engineered cardiac microtissues were maintained at 37°C in a humidified 5% CO_2_ incubator on an orbital shaker rotating at 60 rpm.

### Cell encapsulation

Procyon devices (Procyon Technologies LLC, AZ, USA) with 3 cm^2^ single chamber were conditioned in 95% v/v ethanol for 10 min, 20% v/v ethanol for 10 min and rinsed in sterile phosphate buffered saline in sequential order. The conditioned Procyon devices were loaded with either human Cymerus MSCs (passage 6-7) suspended in 50 μL of Cymerus MSC serum-free expansion medium or 50 μL of Cymerus MSC serum-free expansion medium (as control) using a Hamilton syringe. The cell loading port was then sealed with medical grade adhesive silicone, type A (Nusil, CA, USA) and the excess port was trimmed off. The device was then placed in Cymerus MSC serum-free expansion medium at 37°C in a humidified 5% CO_2_ incubator for 1-3 days before implantation.

### Myocardial ischaemia-reperfusion injury (*in vivo*)

All experimental procedures were approved by the Animal Ethics Committee of St Vincent’s Hospital and were conducted in accordance with the Australian National Health and Medical Research Council guidelines for the care and use of laboratory animals (AEC No. 011/22). All animal procedures conformed to the guidelines from Directive 2010/63/EU of the European Parliament on the protection of animals used for scientific purposes or the NIH guidelines.

#### Part 1 dose-response study

Adult male Sprague Dawley rats aged 8-11 weeks were pretreated with antibiotics (enrofloxacin 175 mg/L, via drinking water) for 3 days prior to surgery. On day 0, anaesthetised animals (Alfaxan, 10 mL/kg, intravenous injection) were intubated and maintained on 2% isoflurane anaesthesia in oxygen at 70-75 breaths per minute. A left anterior thoracotomy was performed at the 4^th^ intercostal space on the left-hand side and the left anterior descending coronary artery was ligated approximately 3 mm below the left atrium using a double knot 6-0 prolene monofilament polypropylene suture tied around a 20mm length of PE100 tubing. Prior to occlusion, lignocaine (12 mg/kg, subcutaneous injection) was administered to reduce the incidence of arrhythmias. Successful coronary artery occlusion was indicated by visible blanching of the myocardium distal to the coronary ligation site and changes in electrocardiogram. Following 45 minutes occlusion, the PE tubing and suture were removed, the thorax and muscle layers were closed with 4-0 silk sutures. Procyon device containing either Cymerus MSCs (1×10^6^, 2×10^6^, 4×10^6^ cells/device) or serum-free expansion medium (as control) was then subcutaneously placed on the dorsal side of the animals. A pocket was created under the skin at the level of the shoulder blades for device insertion, followed by closure of the skin wound. On obtaining spontaneous respiration, a single dose of carprofen (5 mg/kg, subcutaneous injection) was administered for analgesia. Sham control underwent similar open chest surgery without coronary artery occlusion.

#### Part 2 validation study

Male and female Sprague Dawley rats, aged 51-53 weeks, were placed on a high fat rodent diet (total digestible energy: 60.8% lipids, 18.4% protein, and 88.9 g/Kg sucrose; Specialty Feeds; SF13-092). After four weeks, the rats underwent either myocardial ischaemia-reperfusion injury or sham operation (as described above), while continuing the high fat diet for 12 weeks. At the time of reperfusion, a Procyon device containing either 4×10^6^ Cymerus MSCs or serum-free expansion medium (control) was subcutaneously implanted on the dorsal side of the animals.

### Simulated IRI (*in vitro*)

Engineered cardiac microtissues were subjected to 3 hours of hypoxia and 24 hours of reoxygenation to simulate IRI. Hypoxia was induced in a hypoxic chamber (STEMCELL Technologies) where oxygen was purged by pure nitrogen gas for 15 minutes and using a buffer simulating the conditions of ischaemia (in mmol/L: 1.0 KH_2_PO_4_, 10.0 NaHCO_3_, 1.2 MgCl_2_.6H_2_0, 25.0 Na(4-(2-hydroxyethyl)-1-piperazineethanesulfonic acid) (HEPES), 74.0 NaCl, 16.0 KCl, 1.2 CaCl_2_ and 10 mM 2-deoxyglucose, pH 6.7), gassed with pure nitrogen gas for 5 minutes. Reoxygenation was achieved by replacing the buffer with spheroid medium and culturing in a humidified incubator at 37°C (∼21% O_2_). Engineered cardiac microtissues cultured in spheroid medium at 37°C in a humidified CO_2_ incubator throughout the hypoxia and reoxygenation period was served as the normoxic control group. Engineered cardiac microtissues were randomly assigned to receive concentrated Cymerus MSC serum-free expansion medium (as vehicle control) or concentrated conditioned media of Cymerus MSCs at reoxygenation for 24 hours.

### Mass spectrometry-based proteomics

For proteomic analysis of encapsulated Cymerus MSCs (cell lysate) and their secretome (conditioned media, CM), samples were solubilized in 1% (v/v) sodium dodecyl sulphate (SDS) containing 50 mM HEPES (pH 8.0) and HALT protease and phosphatase inhibitor (#78442, Thermo Fisher Scientific). Further, global cell proteomics was performed on a monolayer of Cymerus MSCs (2D) and human iPSCs (iPS-Foreskin-2 cell line) for comparative purpose. Protein lysates were further homogenized by tip-probe sonication on ice and quantified by microBCA (Thermo Fisher Scientific) ^14^.

Cell and secretome samples (10 µg protein) were reduced and alkylated with 10 mM DTT and 20 mM IAA as described^37^. Briefly, samples were prepared using SP3 protocol ^15^ and digested using Trypsin and Lys-C (1:50 and 1:100 enzyme-to-protein ratio, respectively) overnight at 37°C. Samples were acidified after digestion to final concentration of 2% formic acid before vacuum lyophilisation. Samples were reconstituted in 10 µL of 0.07% (v/v) trifluoroacetic acid in LC-MS grade water, peptides quantified using fluorometric peptide assay (Thermo Fisher Scientific). LC-MS data acquisition was performed on Q Exactive HF-X benchtop Orbitrap mass spectrometer coupled with UltiMate™ NCS-3500RS nano-HPLC and operated with Xcalibur software as previously described ^14,16^.

For proteomics analyses, we investigated biological replicates for global cell and secretome proteomics: encapsulated Cymerus MSCs pre-implantation (n=2), encapsulated Cymerus MSCs post-implantation (*ex vivo* cultured) (n=3), 2D Cymerus MSCs monolayer (n=2), iPSCs (n=3). MS-based proteomics data is deposited to the ProteomeXchange Consortium via the MassIVE partner repository and available via MassIVE with identifier (MSV000097257).

DIA-MS spectra were processed using DIA-NN software (v1.9) ^17^ as previously reported ^14,18^. The DIA-MS spectra were searched against human proteome database (UP000005640, #83,401). Perseus (v2.0.11) was applied for data processing and analysis, with scatter plots/bar charts generated using GraphPad Prism (v8.0.1) or Microsoft Excel. Protein intensities were log2 transformed and normalized using quantile normalization. Proteins were subjected to PCA and unpaired student’s t-test or Welch’s T-test with missing values imputed from normal distribution (width 0.3, downshift 1.8). g:Profiler and Reactome pathway databases were utilized for Gene Ontology functional enrichment and network/pathway analysis, significance P < 0.05.

### Statistics

Data are expressed as mean ± standard error of the mean (SEM) with three significant figures. Significance of the differences was evaluated using Student’s or Welch’s t-test, one-way or two-way ANOVA followed by Tukey’s multiple comparison post hoc analysis where appropriate. P < 0.05 is considered statistically significant.

## Results

### Subcutaneously implanted Cymerus MSCs in Procyon device protects young rats from reperfused myocardial infarction

To assess the therapeutic effect of Cymerus MSCs on myocardial infarction, the Procyon device containing Cymerus MSCs was subcutaneously implanted at reperfusion after coronary artery ligation. Pre-operative echocardiography showed no differences in cardiac function among the groups. Following reperfused myocardial infarction, immunocompetent young rats exhibited a progressive deterioration in cardiac function over the 12-week reperfusion period (Figure 1A). However, treatment with Cymerus MSCs encapsulated within the Procyon immunoisolation device significantly preserved cardiac function, showing a significant improvement in ejection fraction at a dose of 4×10^6^ cells by 12 weeks post-infarction (59.9 ± 0.853% versus 49.3 ± 1.98% in control, P < 0.05, Figure 1B). The improvement in cardiac function was associated with reductions in cardiac hypertrophy (Figure 1C), cardiomyocyte hypertrophy (Figure 1D), infarct scar size (Figure 1E), and interstitial fibrosis (Figure 1F), while the myocardial vascular density was similar among groups (lectin^+^/mm^2^ = 2470 ±77.7 in Sham, 2140 ± 60.3 in Control, 2310 ± 81.7 in 1×10^6^ Cymerus MSCs, 2190 ± 150 in 2×10^6^ Cymerus MSCs, and 2270 ± 44.1 in 4×10^6^ Cymerus MSCs; P > 0.05, n=4-6).

**Figure 1.**
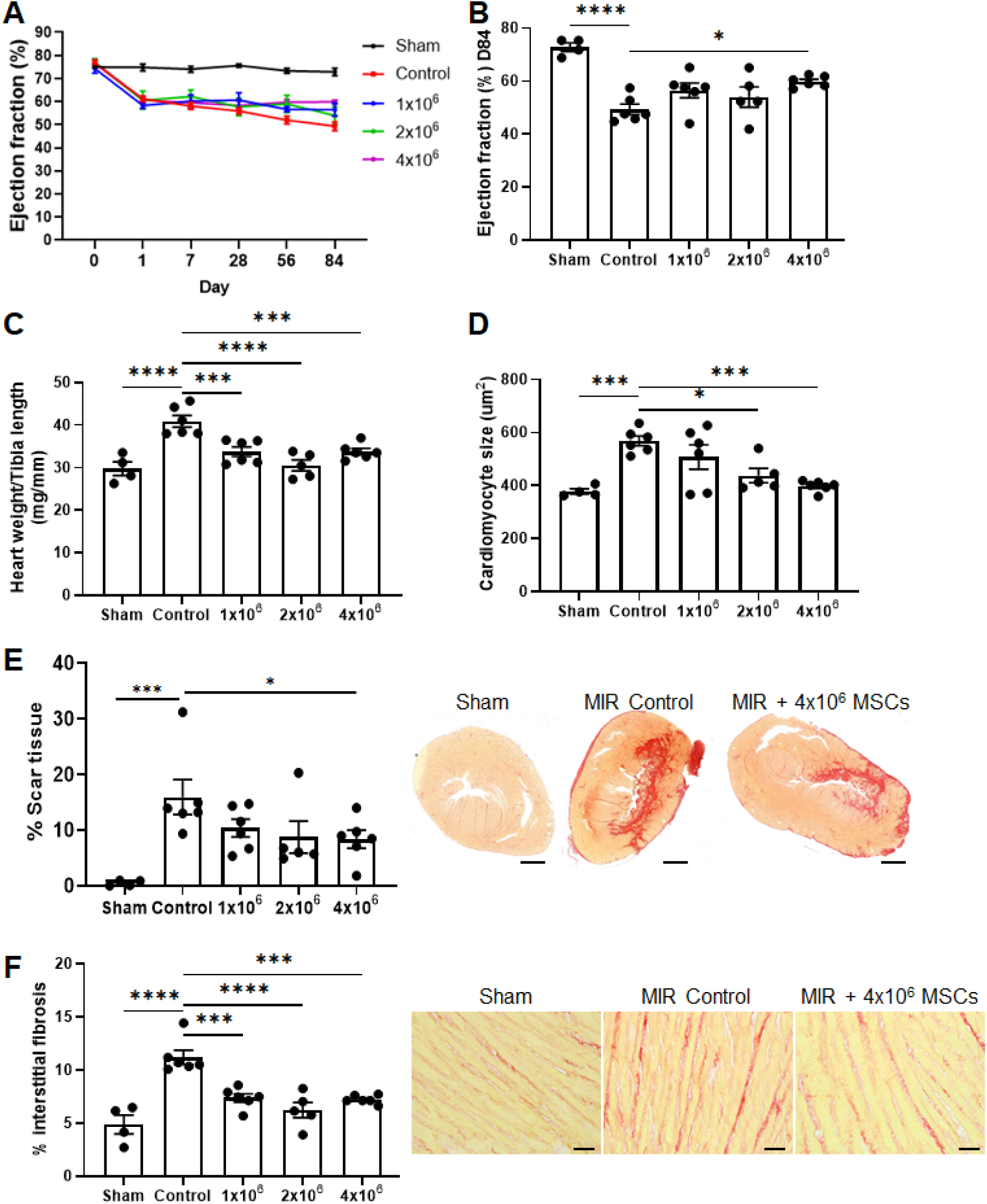
Cardioprotective effects of Cymerus MSCs encapsulated within a Procyon immunoisolation device in young rats following reperfused myocardial infarction. (A) Changes in the left ventricular ejection fraction from baseline over a 12-weeks of reperfusion period. (B) Changes in the left ventricular ejection fraction on day 84 (D84) following reperfused myocardial infarction. (C-F) Cardiac structural and histological assessments on day 84 following reperfused myocardial infarction: (C) normalised heart weight, (D) relative cross-sectional area of cardiomyocytes, (E) infarct scar size expressed as the percentage of fibrotic scar area over total left ventricle and representative cross-sectional images of myocardium stained with picrosirius red (scale bar = 2 mm), and (F) percentage of interstitial fibrosis in the remote myocardium and representative cross-sectional images of myocardium stained with picrosirius red (scale bar = 50 µm). n = 4-6 rats. Data are presented as mean ± SEM. *P < 0.05, ***P < 0.001, ****P < 0.0001 by one-way ANOVA with Bonferroni post hoc test.

### Cardioprotective efficacy in aged and metabolically stressed male and female rats

To further validate the cardioprotective effects of Cymerus MSCs in a clinically relevant model, immunocompetent, middle-aged male and female Sprague-Dawley rats were utilized. These rats were fed a high-fat diet and subjected to reperfused myocardial infarction to mimic an AMI patient with common comorbidities of aging and metabolic stress. At the start of the study (ages 50-52 weeks), female rats exhibited lower body weights than male rats, a difference that remained consistent throughout the study period. Over the 108-day of high-fat diet feeding, all rats showed weight gain regardless of sex, with no significant differences between treatment groups. While female rats consistently maintained lower overall body weights compared to males, the rate of weight gain was similar across all groups (Table 1).

**Table 1.** Body weight and blood biochemical values of aged rats.

| Experimental groups | Male |  |  | Female |  |  |
| --- | --- | --- | --- | --- | --- | --- |
|  | Sham | IR | IR+Cymerus | Sham | IR | IR+Cymerus |
| Sample size | 6 | 7 | 7 | 7 | 7 | 7 |
| Body weight (g), day -28 (before commencing HFD) | 680 ± 49.5 | 677 ± 32.7 | 711 ± 26.2 | 441 ± 21.6 | 406 ± 22.5 | 396 ± 28.6 |
| Body weight (g), day 0 | 775 ± 49.3 | 757 ± 39.0 | 775 ± 26.0 | 502 ± 24.4 | 459 ± 33.0 | 453 ± 36.4 |
| Body weight (g), day 28 | 772 ± 49.1 | 744 ± 33.7 | 758 ± 25.5 | 497 ± 26.1 | 442 ± 31.0 | 425 ± 33.9 |
| Body weight (g), day 56 | 794 ± 45.9 | 770 ± 37.3 | 799 ± 29.4 | 524 ± 21.8 | 478 ± 38.7 | 458 ± 39.6 |
| Body weight (g), day 84 | 804 ± 49.3 | 789 ± 40.8 | 819 ± 29.1 | 554 ± 22.5 | 496 ± 47.3 | 497 ± 48.0 |
| Blood glucose (mmol/L), day -28 (before commencing HFD) | 6.42 ± 0.36 | 6.74 ± 0.18 | 6.80 ± 0.50 | 6.11 ± 0.18 | 6.93 ± 0.32 | 7.39 ± 0.19 |
| Blood glucose (mmol/L), day 0 | 7.73 ± 0.41 | 8.64 ± 0.49 | 8.50 ± 0.43 | 7.47 ± 0.46 | 7.79 ± 0.19 | 7.64 ± 0.23 |
| Blood glucose (mmol/L), day 28 | 9.22 ± 0.29 | 9.06 ± 0.48 | 8.93 ± 0.47 | 9.24 ± 0.45 | 8.04 ± 0.32 | 8.59 ± 0.42 |
| Blood glucose (mmol/L), day 56 | 8.42 ± 0.17 | 8.41 ± 0.31 | 9.19 ± 0.81 | 9.07 ± 0.22 | 8.77 ± 0.25 | 8.55 ± 0.52 |
| Blood glucose (mmol/L), day 84 | 9.97 ± 0.45 | 9.24 ± 0.84 | 10.77 ± 0.83 | 9.06 ± 0.62 | 9.89 ± 0.43 | 8.86 ± 0.51<sup>#</sup> |
| †HbA1C (%) | 3.96 ± 0.11 | 3.92 ± 0.13 | 3.95 ± 0.08 | 3.87 ± 0.10 | 3.55 ± 0.07 | 3.75 ± 0.10 |
| †Total cholesterol (mmol/L) | 2.72 ± 0.29 | 2.57 ± 0.12 | 2.47 ± 0.18 | 2.18 ± 0.17 | 1.88 ± 0.18 | 1.94 ± 0.10 |
| †Triglyceride (mmol/L) | 1.70 ± 0.15 | 1.64 ± 0.16 | 2.14 ± 0.22 | 2.22 ± 0.31 | 2.27 ± 0.41 | 1.81 ± 0.25 |
| †HDL (mmol/L) | 1.76 ± 0.16 | 1.63 ± 0.08 | 1.61 ± 0.15 | 1.65 ± 0.11 | 1.47 ± 0.10 | 1.48 ± 0.09 |
| †LDL (mmol/L) | 0.65 ± 0.12 | 0.51 ± 0.04 | 0.45 ± 0.08 | 0.17 ± 0.02<sup>####</sup> | 0.13 ± 0.02<sup>##</sup> | 0.13 ± 0.01<sup>##</sup> |
| †CRP (pg/mL) | 1280 ± 203 | 2630 ± 898 | 1440 ± 228 | 2220 ± 666 | 5700 ± 1960 | 677 ± 102<sup>**</sup> |
| †IL-6 (pg/mL) | 500 ± 263 | 398 ± 175 | 180 ± 59.8 | 611 ± 207 | 395 ± 62.9 | 152 ± 34.2 |
| †IL-1β (pg/mL) | 62.5 ± 18.7 | 69.1 ± 24.6 | 66.5 ± 35.2 | 67.7 ± 14.8 | 63.2 ± 11.8 | 44.2 ± 9.16 |
| †TNFα (pg/mL) | 15.6 ± 4.34 | 18.2 ± 5.49 | 17.9 ± 8.80 | 19.6 ± 4.38 | 17.6 ± 3.06 | 12.0 ± 2.03 |

†Measurements were performed on day 84 endpoint. HFD (high fat diet), HDL (high-density lipoprotein), LDL (low-density lipoprotein), CRP (c-reactive protein), IL (interleukin), TNFα (tumor necrosis factor-alpha).
<sup>#</sup>P<0.05, <sup>##</sup>P<0.01, <sup>####</sup>P<0.0001 versus male cohort of the same treatment group. <sup>\*\*</sup>P<0.01 versus female IR group.

Blood glucose concentrations were elevated in both male and female rats after 108 days on the high-fat diet, with no significant differences between sexes or treatment groups, except that female rats treated with Cymerus MSCs had lower blood glucose concentrations than their male counterparts (Table 1). Other blood biochemical markers, such as total cholesterol, triglyceride and high-density lipoprotein, were generally comparable across sexes and groups, except for low-density lipoprotein levels, which were significantly lower in female rats than in males (Table 1). This model effectively simulates metabolic stress, and comorbidity risks common in middle-aged AMI patients, providing a clinically relevant preclinical framework for assessing the cardioprotective efficacy of Cymerus MSCs.

Following reperfused myocardial infarction, middle-aged male and female rats displayed a progressive decline in cardiac function over the 12-week reperfusion period (Figure 2A). However, treatment with Cymerus MSCs encapsulated within the Procyon immunoisolation device significantly preserved cardiac function in this pre-clinical animal model. By 12 weeks post-infarction, both male and female rats showed significant improvement in ejection fraction (Figure 2A-B), with effects similar to those observed in younger rats (Figure 1). This improvement was accompanied by significant attenuation of adverse cardiac remodelling, including reductions in cardiac hypertrophy (Figure 2C), cardiomyocyte hypertrophy (Figure 2D, Supplementary Figure S1), infarct scar size (Figure 2E, Supplementary Figure S1), and interstitial fibrosis (Figure 2F, Supplementary Figure S1). Similar to younger male rats, myocardial vascular density in the remote myocardium was comparable among groups in middle-aged male rats (2770 ± 214 lectin^+^/mm^2^ in Sham, 2450 ± 86.8 lectin^+^/mm^2^ in Control, and 2910 ± 130 lectin^+^/mm^2^ in Cymerus MSCs, P > 0.05, n=6-7). Interestingly, in middle-aged female rats, myocardial vascular density was significantly higher in the Cymerus MSC-treated group compared to the control group (3060 ± 183 lectin^+^/mm^2^ in Cymerus MSCs versus 2410 ± 158 lectin^+^/mm^2^ in Control, P < 0.05, n=7). Analysis of inflammatory cytokine levels at the 12-week post-myocardial infarction endpoint showed an increase in the inflammatory marker CRP, though it did not reach statistical significance. Treatment with Cymerus MSCs reduced CRP levels, achieving statistical significance in female rats. Additionally, Cymerus MSCs reduced inflammatory cytokines IL-6 and TNFα, with a stronger effect observed in female rats (Table 1). These results underscore the robust therapeutic potential of Cymerus MSC treatment across age groups and sexes.

**Figure 2.**
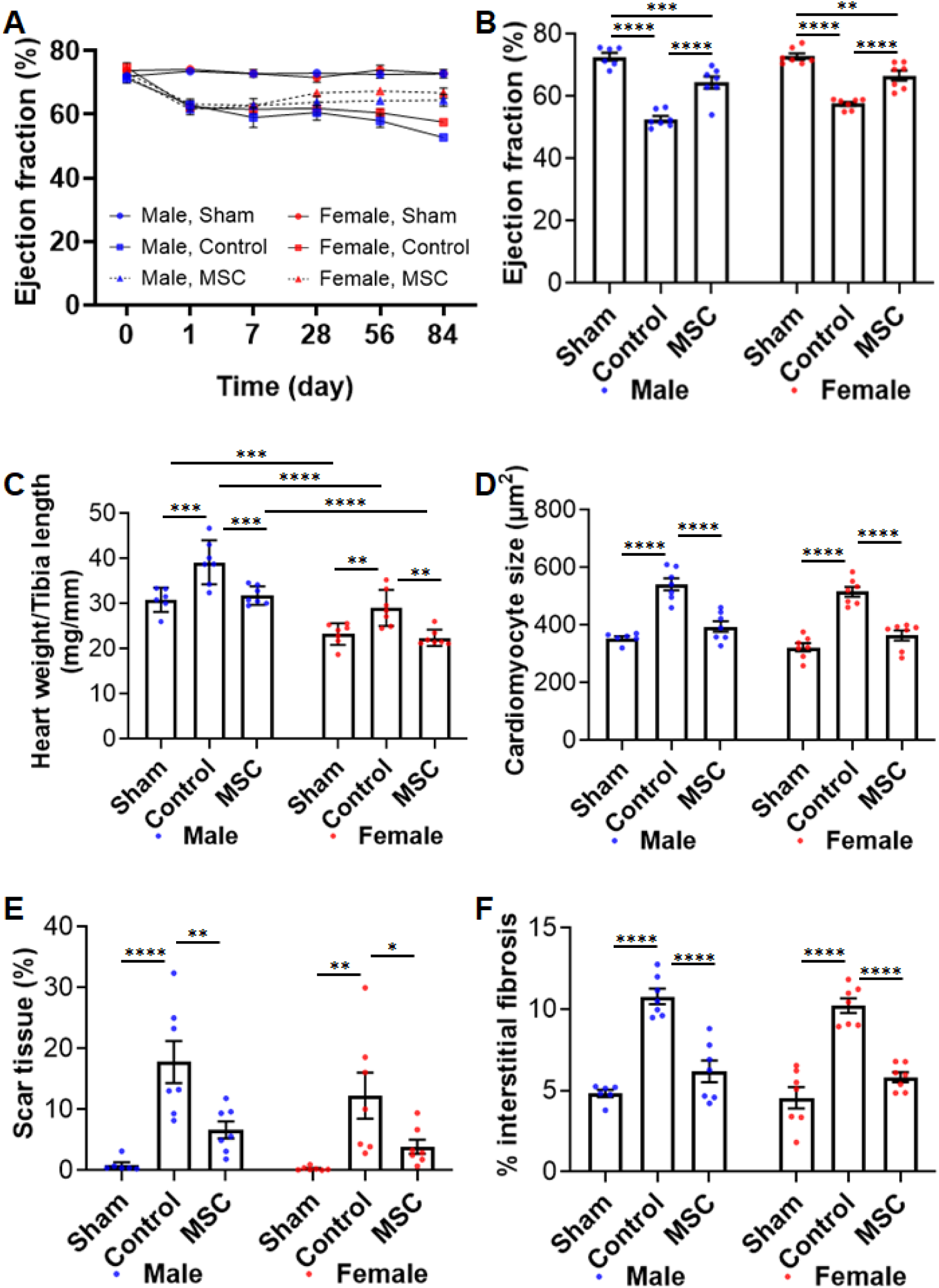
Cardioprotective effects of Cymerus MSCs encapsulated within a Procyon immunoisolation device in middle-aged male and female rats following reperfused myocardial infarction. (A) Changes in the left ventricular ejection fraction from baseline over a 12-weeks of reperfusion period. (B) Changes in the left ventricular ejection fraction on day 84 following reperfused myocardial infarction. (C-F) Cardiac structural and histological assessments on day 84 following reperfused myocardial infarction: normalised heart weight (C), relative cross-sectional area of cardiomyocytes in the remote myocardium (D), infarct scar size expressed as the percentage of fibrotic scar area over total left ventricle (E), and percentage of interstitial fibrosis in the remote myocardium (F). n = 6-7 rats. Data are presented as mean ± SEM. *P < 0.05, **P < 0.01, ***P < 0.001, ****P < 0.0001 by one-way ANOVA with Bonferroni post hoc test across both sexes.

### Post-implantation morphological adaptations of Cymerus MSCs

To evaluate the effects of *in vivo* implantation on Cymerus MSCs, we performed histological analysis on Procyon devices retrieved 12 weeks after subcutaneous implantation in immunocompetent rats. Human-specific KU80 staining confirmed that encapsulated Cymerus MSCs remained fully contained within the inner membranes of the devices (Supplementary Figure S2). The encapsulated cells appeared viable, exhibiting well-preserved morphology with intact membranes and clearly defined nuclei, and showed no signs of apoptosis, as indicated by the absence of cleaved caspase-3 (Figure 3A-B, Supplementary Figure S2).

**Figure 3.**
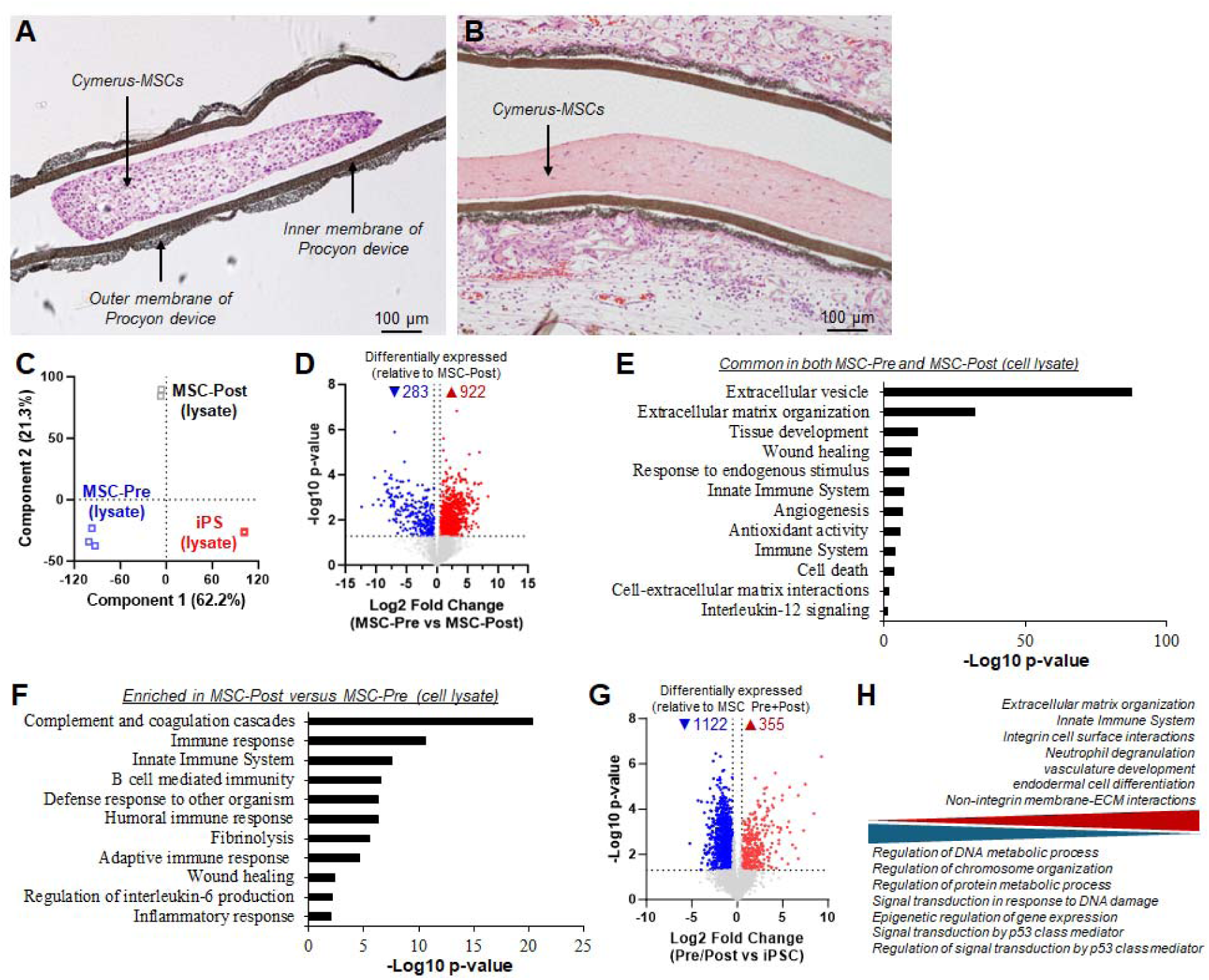
Morphological and proteomic network changes in encapsulated Cymerus MSCs following *in vivo* implantation in rats subjected to reperfused myocardial infarction. (A-B) Haematoxylin and eosin stained cell encapsulated Procyon device at 0 (**A**, Pre) and 12 (**B**, Post) weeks post-implantation. (**C-H**) Global proteome analysis of encapsulated Cymerus MSCs at 0 (MSC-Pre) and 12 (MSC-Post) weeks post-implantation, and human iPSCs. (**C**) Principal component analysis of indicated cellular proteomes. (**D**) Volcano plot of cellular proteome of MSC- Pre relative to MSC-Post, based on Welch’s T-Test, P < 0.05 and log2 fold change of 0.5. Functional enrichment analysis of Reactome and GO networks/terms enriched (P < 0.05) in proteins commonly identified and expressed in MSC-Pre and MSC-Post cellular proteome (**E**), and in proteins significantly upregulated in expression in MSC-Post cellular proteome relative to MSC-Pre cellular proteome (**F**). (**G**) Volcano plot of cellular proteome of encapsulated Cymerus MSCs (MSC-Pre and MSC-Post combined) relative to iPSCs. Based on Welch’s T-Test, P < 0.05 and log2 fold change of 0.5. (**H**) Trend enrichment analysis of Reactome networks and GO biological processes. Biological process enriched (P < 0.05) in differentially expressed Cymerus MSC (MSC-Pre and MSC-Post combined) relative to iPSC cellular proteome.

Similar to the control device (lacking encapsulated Cymerus MSCs), minimal leukocyte infiltration was observed around the devices containing Cymerus MSCs at 12 weeks post-implantation, suggesting an absence of immunological or xeno-immune reactions toward the foreign material and cells (Figure 3B). Over the 12-week period, the encapsulated cells continued to express stromal/mesenchymal marker vimentin and reorganized into tissue-like structures supported by self-secreted extracellular matrices within the devices (Figure 3A-B, Supplementary Figure S2).

Notably, cartilaginous tissue formation was detected in one of the 28 implanted devices, as evidenced by positive Alcian blue staining (Supplementary Figure S2). Immunostaining with the pan-cell cycle marker Ki67 revealed a reduction in the proliferative activity of Cymerus MSCs at 12 weeks post-implantation compared to pre-implantation (Supplementary Figure S2). Additionally, extensive vascularization was observed along the outer membranes of the devices, with rodent-specific lectin staining indicating integration with the host vasculature (Supplementary Figure S2). These findings suggest that the encapsulated Cymerus MSCs retain structural integrity and viability while forming organized tissue structures within the device. Further, while the proliferative activity of encapsulated MSCs declines over time, and the outer membrane matrix promotes robust vascularization, enhancing host-device integration.

### Proteomic profile of encapsulated Cymerus MSCs following *in vivo* implantation

To better understand the temporal cellular changes in Cymerus MSCs during implantation, a global (untargeted) proteomic analysis was performed on encapsulated Cymerus MSCs both before implantation and 12 weeks post-implantation. Using stringent identification criteria, 3356 proteins were identified in pre-implantation phase, and 1782 proteins in post-implantation (Supplementary Table 1, Supplementary Figure S3A). Principal component analysis revealed distinct clustering of encapsulated Cymerus MSCs, including their relation to human iPSC proteome (Figure 3C, Supplementary Table 1).

#### Conserved proteome following encapsulation

Cymerus MSCs retain a conserved core cellular proteome following encapsulation within the immunoisolation devices. When comparing monolayer cultured MSCs (2D MSC) to encapsulated MSCs pre-implantation (MSC-Pre), minimal impact on the cellular proteomic profile was observed, with 3,357 proteins commonly identified based on abundance/identification (>75% of cellular proteome) (Supplementary Figure S3B-E, Supplementary Table 2). Key MSC markers within this subset include NCAM1, ANPEP, MME, THY1, ALCAM, ITGB1, NT5E, MCAM, PDGFRB, SRC, ENG, and CD44. Of the 475 proteins with altered expression associated with encapsulation, enrichment was observed in biological process ontologies related to cell adhesion (HAS1, LAMC1, LAMB1, CBFB), extracellular matrix organization (HAS1, COL1A2, LAMC1, ERO1A), wound healing (CLIC1, VKORC1, AK3, TOR1A, MYL12A), and collagen metabolic process (COL1A2, CTSB, MMP14, MRC2, FAP) (Supplementary Table 3, Supplementary Figure S3E).

#### Proteomic changes in the cellular landscape post-implantation

To compare cellular proteome landscape of MSC-Pre with encapsulated Cymerus MSCs post-implantation (MSC-Post), a global analysis was performed to identify changes that reflect the proteome environment of the *in vivo* implant. Of the 1547 proteins commonly identified (43.1% of the Cymerus MSC cellular proteome containing 3591 proteins, Supplementary Table 5), key biological processes were enriched, including extracellular vesicle dynamics (RAB11B, RAB15, RAC1, CDC42, ARPC2, TOLLIP, MVP), extracellular matrix organization (LAMA4, COL3A1, MMP2, HAPLN1, COL11A1), tissue development (EGFR, TGFBI, AKT1, FN1, POSTN, MAPK3), angiogenesis (NRP1, AKT1, RRAS, LOXL2), antioxidant activity (PRDX5, PRDX4, GPX7, CP, APOE), response to endogenous stimulus (EGFR, ANXA1, SERPINF1, GSTM3, CAV1), cell death regulation (AKT1, PRKAA1, MAPK3, CLU), immune regulation protein networks (ACTB, HP, ANPEP, RAC2, CD44), membrane trafficking (ACTB, EGFR, MAN1A1, DAB2, COPZ2), interleukin-12 signalling (CDC42, RAP1B, P4HB, MIF, ANXA2), and cell-extracellular matrix interactions (ACTB, LIMS2, LIMS1, ILK, PARVA). Notably, MSC markers CD44 and vimentin (VIM) were also identified (Figure 3E, Supplementary Table 6).

A subset of 179 proteins (5.0% of the total cellular proteome) were uniquely identified in the post-implant cell proteome, associated with immune effector process (IGKV3-15, IGHV3OR16-12, IGHV1OR15-1, IGHV1-45, IGKV6D-21, NEDD4), anti-inflammatory response (LGALS9, A2M, PTPN6, IRF5, IKBKG), complement activation (C4A, C3, F13A1, F2, C1R, PLG), wound healing response (F13A1, F2, PLG, FGA, FGB), and fibrinolysis (FGG, VTN, CPB2) (Figure 3F, Supplementary Table 7). Together, these findings suggest that MSCs undergo proteome reprogramming upon implantation to adapt to the *in vivo* environment, promoting immune modulation, tissue repair, and integration into the surrounding tissue.

To further understand MSC proteome reprogramming relative to their iPSC origin, Welch’s T-Test paired analyses identified 355 conserved MSC-associated proteins in both MSC-Pre and MSC-Post (Figure 3G, Supplementary Table 5). These proteins were enriched in key biological processes, including extracellular matrix organisation, innate immune system, integrin cell surface interactions, collagen formation, and vasculature development (Figure 3H, Supplementary Table 8). In contrast, 1122 proteins were significantly enriched in iPSCs compared to MSCs, primarily associated with cell cycle regulation, DNA metabolism, chromosome organisation, epigenetic regulation of gene expression, and p53-mediated signalling (Figure 3G-H, Supplementary Table 5 and 9). This comparison highlights how MSCs and iPSCs are specialised at the cellular proteome level for their respective biological functions. MSCs are primed to maintain tissue homeostasis and respond to injury or inflammation, while iPSCs maintain pluripotency and the potential to differentiate into various cell types, a key characteristic of stem cell biology.

### Adaptive response of Cymerus MSCs secreting factors implicated in cardiac biology and tissue remodelling post-implantation

Proteome profiling was performed to characterise the secretome composition of Cymerus MSCs and assess temporal changes in their secreted protein profile following *in vivo* implantation (*ex vivo* cultured). A total of 448 secreted proteins were identified (in at least one sample, and 69 proteins in at least two) from MSC-Pre, and 815 proteins (589 proteins in at least two samples) from MSC-Post (Figure 4A, Supplementary Table 10). The encapsulation of Cymerus MSCs minimally affects their secretome when compared to Cymerus MSCs cultured in a monolayer (Supplementary Figure S4). Principal component analysis demonstrated a distinct clustering pattern in the MSC-Post secretome, indicating significant shifts in protein expression post-implantation (Figure 4B).

**Figure 4.**
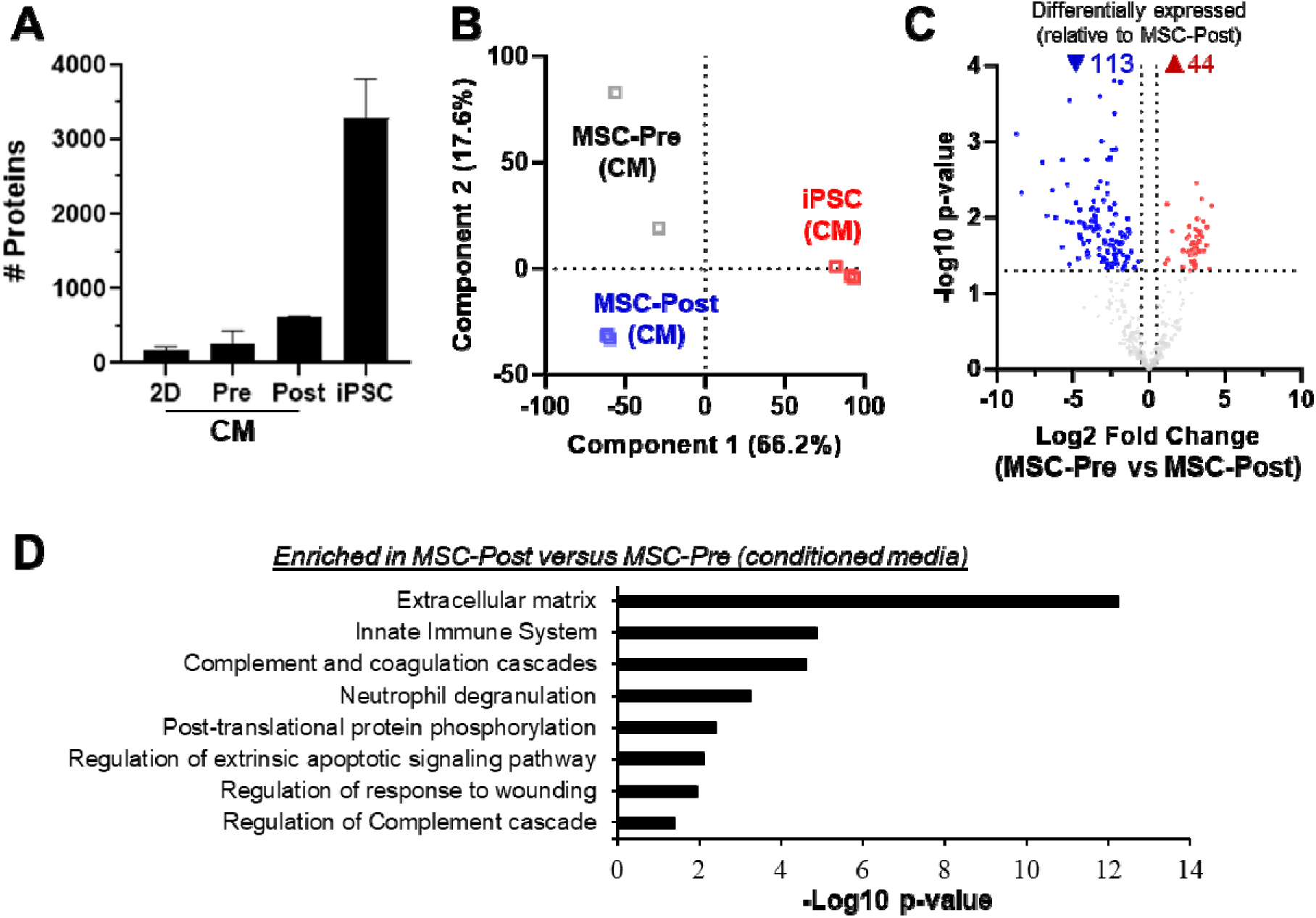
Proteomic network changes in secretome of encapsulated Cymerus MSCs following *in vivo* implantation in rats subjected to reperfused myocardial infarction. (**A**) Quantitative proteomic profiling reveals in-depth coverage and identified protein in conditioned media (CM) collected from 2D Cymerus MSC (2D), encapsulated Cymerus MSCs at 0 (MSC-Pre) and 12 (MSC-Post) weeks post-implantation, and human iPSCs. (**B**) Principal component analysis of indicated conditioned media (secretome) of Cymerus MSCs and human iPSC proteome. (**C**) Volcano plot of conditioned media proteome of MSC-Pre relative to MSC-Post. Based on Welch’s T-Test, P < 0.05 and log2 fold change of 0.5. (**D**) Functional enrichment analysis of Reactome and GO networks/terms enriched (P < 0.05) in proteins significantly upregulated in expression in MSC-Post secretome relative to MSC-Pre secretome.

While MSC markers CD44 and VIM were detected in both MSC-Pre and MSC-Post secretomes (Supplementary Table 11), a greater number of distinct proteins were identified in the secretome from the MSC-Post group compared to the MSC-Pre group (Figure 4C). Functional enrichment analyses of this enriched protein subset revealed pathways associated with the innate immune system (ITIH3, ARHGDIA, ANXA1, SUMO2, PLG), regulation of extrinsic apoptotic signalling pathway (FGG, FGB, SERPINE1), extracellular matrix (ANXA1, MRC2, VCAN), regulation of complement cascade (SUMO2, SYNCRIP, RAD23B), neutrophil degranulation (ANXA1, PLG, LTA4H), and regulation of response to wounding (FGG, FGB, ANXA1, PLG, SERPINE1) (Figure 4D, Supplementary Table 12 and 13). This analysis highlighted an enrichment of secreted proteins in the MSC-Post group (relative to MSC-Pre) associated with wound repair, protection, inflammation, immune regulation, and tissue remodelling.

To further investigate Cymerus MSC secretome reprogramming in relation to encapsulated states (both pre- and post-implantation) and iPSC secreted profiles, a Welch’s T-Test was performed. This identified significant differences between Cymerus MSC and iPSC secretomes, with 58 proteins enriched and 193 downregulated in Cymerus MSC relative to iPSC (Supplementary Table 14). iPSC secretome enrichment includes biological processes and pathway networks associated with RNA metabolism, cap-dependent translation, cell cycle, protein metabolism, RNA degradation, and DNA replication pathways (Supplementary Table 15). In contrast, the Cymerus MSC secretome is enriched in ECM processes, extracellular vesicles, and oxidant detoxification (Supplementary Table 16). These findings suggest that Cymerus MSCs undergo significant reprogramming in response to *in vivo* implantation, with distinct changes in their secretome profile that may contribute to their therapeutic potential.

### Cardioprotective effect of Cymerus MSC secretome in a human engineered cardiac microtissue model of ischaemia-reperfusion injury

To assess the cardioprotective effect of Cymerus MSC secretome in a human context, we generated engineered cardiac microtissues composed of human iPSC-derived cardiomyocytes, endothelial cells, vascular smooth muscle cells, and cardiac fibroblasts. Within these 3D engineered cardiac microtissues, cardiomyocytes self-organised into a ring-like structure, mimicking native cardiac tissue architecture. Endothelial cells localised around the lumen-like inner core, suggesting the formation of primitive vascular structures. Meanwhile, vimentin positive mesenchymal stromal cells/fibroblasts and smooth muscle cells were distributed throughout the engineered cardiac microtissue and distributed among the cardiomyocytes, contributing to the structural complexity and cellular interactions necessary for functional cardiac tissue development (Figure 5A).

**Figure 5.**
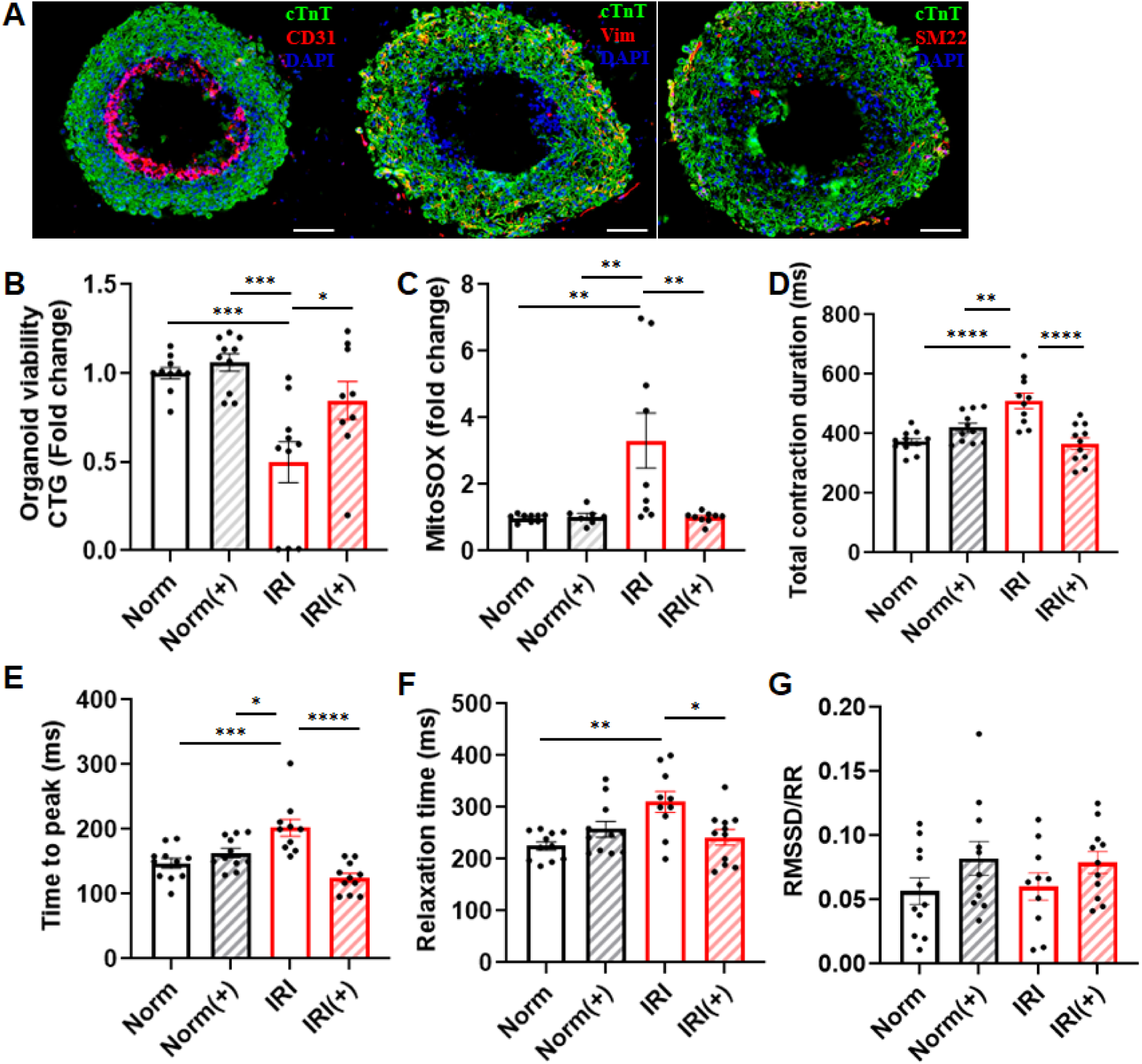
Cardioprotective effects of Cymerus MSC secretome in a pre-clinical human engineered cardiac microtissue model of ischaemia-reperfusion injury. (A) Representative images of engineered cardiac microtissues stained with cardiac troponin T (cTnT, a marker for cardiomyocytes), CD31 (a marker for endothelial cells), SM22 (transgelin, a marker for smooth muscle cells) and vimentin (a marker for mesenchymal stromal cells and fibroblasts). Scale bar = 100 µm. (B) Viability of engineered cardiac microtissues assessed by CellTiter-Glo assay (n=9-10). (C) Mitochondrial ROS production of engineered cardiac microtissues assessed by MitoSOX Red indicator (n=7-9). (D-G) The contraction profile of engineered cardiac microtissues; total contraction duration (D), time-to-peak (E), relaxation time (F), and beat rate variability calculated by the root mean square of successive differences normalised by the R-R interval (G) (n=10-11). Engineered cardiac microtissues were subjected to normoxia (Norm) or simulated IRI conditions (IRI) and treated with vehicle control or 1000 µg of Cymerus MSC conditioned media (+). Data are shown as mean ± SEM. *p<0.05, **p<0.01, ***p<0.001, ****p<0.0001 by one-way ANOVA with Bonferroni post-hoc test.

Engineered cardiac microtissues subjected to simulated IRI showed a significant decrease in viability compared to the normoxia group. Treatment with Cymerus MSC secretome significantly improved the viability of engineered cardiac microtissues when compared to the IRI control group (Figure 5B). Similarly, mitochondrial ROS levels were significantly elevated following simulated IRI but were significantly reduced with Cymerus MSC secretome treatment (Figure 5C). We also assessed the contraction kinetics of the engineered cardiac microtissues, including time-to-peak contraction, relaxation time, and beat rate variability. Compared to the normoxia control, engineered cardiac microtissues exposed to simulated IRI showed a significantly prolonged total contraction duration, time-to-peak, and relaxation time. These changes were effectively prevented by Cymerus MSC secretome treatment (Figure 5D-F). Beat rate variability remained consistent across all experimental groups (Figure 5G). Under normoxic conditions, treatment with Cymerus MSC secretome had no impact on engineered cardiac microtissue viability, mitochondrial ROS production, or contraction kinetics compared to the normoxia control (Figure 5B-G).

## Discussion

This study presents an innovative strategy for secretome-based cardioprotection using human iPSC-derived MSCs delivered via the Procyon immunoisolation device. In an immunocompetent rat model of chronic reperfused myocardial infarction, this approach significantly improved cardiac function, reduced fibrosis, and demonstrated long-term therapeutic benefits. Furthermore, the cardioprotective effect of the MSC secretome remained effective in aged and metabolically stressed animals, comorbidities that typically compromise the efficacy of other cardioprotective strategies such as ischaemic conditioning and remote ischaemic preconditioning ^19–21^. This is particularly significant, as many cardioprotective interventions fail to translate into clinical practice due to the prevalence of elderly patients with multiple comorbidities, a population often resistant to conventional treatments. The resilience of MSC secretome therapy under these challenging conditions presents a promising clinically applicable therapeutic avenue. Although the cardioprotective efficacy of Cymerus MSCs in clinical settings has yet to be determined, our data from human iPSC-derived engineered cardiac microtissues suggest their potential translatability to humans. These findings demonstrate significant protection against ischaemia-reperfusion injury, including restored contractile function, enhanced cell viability, and reduced oxidative stress, underscoring the therapeutic potential of Cymerus MSCs. Nevertheless, further investigation using well-designed clinical trials is essential to establish their safety, effectiveness, and long-term benefits in real-world scenarios.

Meta-analyses of clinical trials consistently report strong safety profiles for MSCs, with minimal adverse events such as immune rejection ^22,23^. Previous clinical trials involving intravenous delivery of Cymerus MSCs showed no tumours or other safety concerns over a two-year follow-up period ^12,24^. Our proteomic analysis of both the encapsulated Cymerus MSCs and their secretome post-implantation revealed no expression of oncogenic protein markers or signs of oncogenic transformation. Laboratory studies further confirmed that the MSC secretome does not promote the growth of human cancer cells in culture (Supplementary Figure S5). Beyond *in vitro* assessments, *ex vivo* post-mortem analysis provides additional evidence supporting the safety profile of this approach. Histomorphological examination of rats implanted with devices containing Cymerus MSCs showed no signs of tissue abnormalities or malignancy, supporting that this intervention does not promote a pro-tumorigenic environment. Together, these findings reinforce the safety profile of Cymerus MSCs, affirming their non-oncogenic properties, immune compatibility, and controlled biological activity. These attributes position them as a safe and viable option for regenerative medicine.

MSCs have gained significant attention in regenerative medicine for their potential to protect the heart in various pathological conditions, including myocardial infarction ^6,25^. The cardioprotective effects of MSCs are largely attributed to their paracrine signalling, where they secrete bioactive factors such as cytokines, growth factors, and extracellular vesicles that mediate their therapeutic actions. Rather than transdifferentiating into cardiomyocytes and directly replacing damaged heart cells, MSCs promote tissue repair by reducing arrhythmic risk, modulating inflammation, preventing apoptosis, limiting fibrosis, and promoting angiogenesis. These actions contribute to myocardial preservation and functional recovery following injury ^4,6,26–28^. The paracrine mechanism of MSCs is evidenced by the observation that conditioned media derived from MSCs or cell extracts can exert cardioprotective effects comparable to those achieved by delivering live MSCs to the heart ^27–31^. Our data further supports this paracrine effect, demonstrating cardioprotection when MSCs are delivered via an immunoisolation device implanted remotely from the myocardium ^11^; suggesting that MSCs do not need to be in direct contact with the heart to exert their beneficial effects. A recent study transplanting noncontractile cardiomyocytes further highlights the role of secretome-based cardioprotection, demonstrating that the cardiac benefits are primarily due to the secretion of paracrine factors, rather than cardiomyocyte graft contraction. In this study, engineered noncontractile human iPSC-derived cardiomyocytes, created by knocking out TNNI1 and TNNI3, were able to preserve cardiac function to a similar extent as contractile cardiomyocytes ^32^. Interestingly, some studies have shown that transplanting dead or apoptotic stem cells into the heart can still provide cardioprotection ^33^. This paradox is explained by the “dying stem cell hypothesis,” which suggests that the effect is due to the stimulation of anti-inflammatory cytokines by the dying cells. These cytokines may help limit scar formation and trigger endogenous repair mechanisms ^34^. This finding highlights the resilience of the MSC-mediated paracrine effect, where beneficial signals persist even after cell death, continuing to protect the myocardium.

Optimising the therapeutic potential of Cymerus MSCs in cardiovascular disease requires a thorough understanding of their cardioprotective mechanisms; particularly their interaction with the host microenvironment. Their ability to sense ischaemic cues and secrete bioactive factors is central to their function, yet the precise nature of these interactions in pathological settings remains unclear. Investigating how transplanted MSCs adapt and modulate their secretome in response to host tissue signals will be key to refining their clinical application. Traditional delivery methods, such as intramyocardial or intravenous injection, present challenges, including the inability to retrieve transplanted cells and the difficulty of long-term tracking, as they are cleared from the host body within days ^35–37^, making it challenging to study their *in vivo* behaviour. The present study addresses this challenge by utilising encapsulated Cymerus MSCs, which can be retrieved post-implantation for detailed analysis. This approach provides significant insights into the dynamic reprogramming of MSCs in response to ischaemic injury, highlighting their substantial plasticity and therapeutic potential.

Following *in vivo* implantation, Cymerus MSCs showed significant proteomic shifts, with a significant increase in secreted proteins (589 post-implantation versus 69 pre-implantation) and distinct clustering patterns in principal component analysis. These changes enriched pathways involved in wound repair, immune regulation, and tissue remodelling. Additionally, post-implantation, Cymerus MSCs demonstrated robust molecular reprogramming with 179 unique proteins linked to immune response, anti-inflammatory response, complement activation and wound healing. Their secretory profile shifted from pro-regenerative to immunomodulatory, in line with the inflammatory licensing process described by Hodgson-Garms and colleagues ^38^. In this process, the resting MSCs primarily focus on tissue homeostasis and repair, promoting extracellular matrix deposition and vascular development, with their secretomes rich in extracellular matrix and pro-regenerative proteins ^4,38,39^. Under inflammatory conditions, such as myocardial infarction, these functions are downregulated, and MSCs prioritise the secretion of chemotactic and immunomodulatory proteins ^38,40^. Similarly, preconditioning with pro-inflammatory cytokines enhances the anti-inflammatory and immunomodulatory properties of MSCs ^41^.

The reduced inflammatory markers observed in plasma of rats treated with Cymerus MSCs, coupled with enhanced secretion of immunomodulatory factors ^38,39^, further underscores their therapeutic potential. These results emphasise the ability of Cymerus MSCs to modulate their secretome in response to environmental cues, providing new insights into their efficacy as regenerative therapies, particularly for cardiovascular diseases.

In conclusion, MSCs primarily exert their cardioprotective effects through paracrine signalling, with their secretome driving therapeutic benefits. This shift supports secretome-based therapy as a less invasive and more efficient alternative to traditional direct cell transplantation. Despite extensive research, stem cell therapy for MI has yielded suboptimal outcomes due to the complex pathology of cardiovascular disease and poor cell retention. However, paracrine signalling is now widely recognized as the key mechanism behind MSC-driven cardiac repair. Our study highlights the advantages of encapsulated Cymerus MSCs, which remain viable post-implantation and continuously release therapeutic factors. This approach addresses key translational challenges, including histocompatibility concerns and poor engraftment, making MSC-secretome therapy a scalable and clinically viable strategy for long-term cardioprotection. Furthermore, this study provides valuable insights into the cellular reprogramming of Cymerus MSCs following *in vivo* implantation, revealing conserved and dynamic changes in their proteome that enhance their therapeutic potential for cardiovascular diseases. By deepening our understanding of the cellular and molecular mechanisms driving MSC-based therapies, this work paves the way for more effective and targeted clinical applications in regenerative medicine.

## Statements and Declarations

### Author contributions

SYL conceived the project, designed, and performed the experiments, and analysed and interpreted data. AMK, DWG, JGL, AK, JC, AN, JP, SN, YD, LM, CK, TL, SYL planned and performed experiments, and analysed data. DWG, RHR, JM, KKP, KK, DH, TL, and SYL provided materials and methodologic input. DWG, JGL, JC, JRTD, TH, AN, JJHC, RHR, JLM, KKP, KK, DJH, TL, and SYL provided interpretative review of study findings. All authors participated in manuscript preparation and approved the final version of the manuscript.

## References

1. Wang Y, Li Q, Bi L, Wang B, Lv T, Zhang P. Global trends in the burden of ischemic heart disease based on the global burden of disease study 2021: the role of metabolic risk factors. BMC Public Health. 2025;25:310. doi: 10.1186/s12889-025-21588-9

2. Swaroop G. Post-myocardial Infarction Heart Failure: A Review on Management of Drug Therapies. Cureus. 2022;14:e25745. doi: 10.7759/cureus.25745

3. Tompkins BA, Balkan W, Winkler J, Gyongyosi M, Goliasch G, Fernandez-Aviles F, Hare JM. Preclinical Studies of Stem Cell Therapy for Heart Disease. Circ Res. 2018;122:1006–1020. doi: 10.1161/CIRCRESAHA.117.312486

4. Thavapalachandran S, Le TYL, Romanazzo S, Rashid FN, Ogawa M, Kilian KA, Brown P, Pouliopoulos J, Barry AM, Fahmy P, et al. Pluripotent stem cell-derived mesenchymal stromal cells improve cardiac function and vascularity after myocardial infarction. Cytotherapy. 2021;23:1074–1084. doi: 10.1016/j.jcyt.2021.07.016

5. Kawasumi R, Kawamura T, Yamashita K, Tominaga Y, Harada A, Ito E, Takeda M, Kita S, Shimomura I, Miyagawa S. Systemic administration of induced pluripotent stem cell-derived mesenchymal stem cells improves cardiac function through extracellular vesicle-mediated tissue repair in a rat model of ischemic cardiomyopathy. Regen Ther. 2025;28:253–261. doi: 10.1016/j.reth.2024.12.008

6. Han Y, Yang J, Fang J, Zhou Y, Candi E, Wang J, Hua D, Shao C, Shi Y. The secretion profile of mesenchymal stem cells and potential applications in treating human diseases. Signal Transduct Target Ther. 2022;7:92. doi: 10.1038/s41392-022-00932-0

7. Guo Y, Yu Y, Hu S, Chen Y, Shen Z. The therapeutic potential of mesenchymal stem cells for cardiovascular diseases. Cell Death Dis. 2020;11:349. doi: 10.1038/s41419-020-2542-9

8. Menasché P, Renault NK, Hagège A, Puscas T, Bellamy V, Humbert C, Le L, Blons H, Granier C, Benhamouda N, et al. First-in-man use of a cardiovascular cell-derived secretome in heart failure. Case report. eBioMedicine. 2024;103. doi: 10.1016/j.ebiom.2024.105145

9. Golpanian S, Schulman IH, Ebert RF, Heldman AW, DiFede DL, Yang PC, Wu JC, Bolli R, Perin EC, Moye L, et al. Concise Review: Review and Perspective of Cell Dosage and Routes of Administration From Preclinical and Clinical Studies of Stem Cell Therapy for Heart Disease. Stem Cells Transl Med. 2016;5:186–191. doi: 10.5966/sctm.2015-0101

10. Zhang J, Bolli R, Garry DJ, Marban E, Menasche P, Zimmermann WH, Kamp TJ, Wu JC, Dzau VJ. Basic and Translational Research in Cardiac Repair and Regeneration: JACC State-of-the-Art Review. J Am Coll Cardiol. 2021;78:2092–2105. doi: 10.1016/j.jacc.2021.09.019

11. Kompa AR, Greening DW, Kong AM, McMillan PJ, Fang H, Saxena R, Wong RCB, Lees JG, Sivakumaran P, Newcomb AE, et al. Sustained subcutaneous delivery of secretome of human cardiac stem cells promotes cardiac repair following myocardial infarction. Cardiovasc Res. 2021;117:918–929. doi: 10.1093/cvr/cvaa088

12. Bloor AJC, Patel A, Griffin JE, Gilleece MH, Radia R, Yeung DT, Drier D, Larson LS, Uenishi GI, Hei D, et al. Production, safety and efficacy of iPSC-derived mesenchymal stromal cells in acute steroid-resistant graft versus host disease: a phase I, multicenter, open-label, dose-escalation study. Nat Med. 2020;26:1720–1725. doi: 10.1038/s41591-020-1050-x

13. Yu J, Vodyanik MA, Smuga-Otto K, Antosiewicz-Bourget J, Frane JL, Tian S, Nie J, Jonsdottir GA, Ruotti V, Stewart R, et al. Induced pluripotent stem cell lines derived from human somatic cells. Science. 2007;318:1917–1920. doi: 10.1126/science.1151526

14. Cross J, Rai A, Fang H, Claridge B, Greening DW. Rapid and in-depth proteomic profiling of small extracellular vesicles for ultralow samples. Proteomics. 2024;24:e2300211. doi: 10.1002/pmic.202300211

15. Hughes CS, Moggridge S, Muller T, Sorensen PH, Morin GB, Krijgsveld J. Single-pot, solid-phase-enhanced sample preparation for proteomics experiments. Nat Protoc. 2019;14:68–85. doi: 10.1038/s41596-018-0082-x

16. Tham YK, Bernardo BC, Claridge B, Yildiz GS, Woon LM, Bond S, Fang H, Ooi JYY, Matsumoto A, Luo J, et al. Estrogen receptor alpha deficiency in cardiomyocytes reprograms the heart-derived extracellular vesicle proteome and induces obesity in female mice. Nat Cardiovasc Res. 2023;2:268–289. doi: 10.1038/s44161-023-00223-z

17. Demichev V, Messner CB, Vernardis SI, Lilley KS, Ralser M. DIA-NN: neural networks and interference correction enable deep proteome coverage in high throughput. Nat Methods. 2020;17:41–44. doi: 10.1038/s41592-019-0638-x

18. Fang H, Greening DW. An Optimized Data-Independent Acquisition Strategy for Comprehensive Analysis of Human Plasma Proteome. Methods Mol Biol. 2023;2628:93–107. doi: 10.1007/978-1-0716-2978-9_7

19. Ferdinandy P, Andreadou I, Baxter GF, Botker HE, Davidson SM, Dobrev D, Gersh BJ, Heusch G, Lecour S, Ruiz-Meana M, et al. Interaction of Cardiovascular Nonmodifiable Risk Factors, Comorbidities and Comedications With Ischemia/Reperfusion Injury and Cardioprotection by Pharmacological Treatments and Ischemic Conditioning. Pharmacol Rev. 2023;75:159–216. doi: 10.1124/pharmrev.121.000348

20. Behmenburg F, Heinen A, Bruch LV, Hollmann MW, Huhn R. Cardioprotection by Remote Ischemic Preconditioning is Blocked in the Aged Rat Heart in Vivo. J Cardiothorac Vasc Anesth. 2017;31:1223–1226. doi: 10.1053/j.jvca.2016.07.005

21. Boengler K, Buechert A, Heinen Y, Roeskes C, Hilfiker-Kleiner D, Heusch G, Schulz R. Cardioprotection by ischemic postconditioning is lost in aged and STAT3-deficient mice. Circ Res. 2008;102:131–135. doi: 10.1161/CIRCRESAHA.107.164699

22. Wang Y, Yi H, Song Y. The safety of MSC therapy over the past 15 years: a meta-analysis. Stem Cell Res Ther. 2021;12:545. doi: 10.1186/s13287-021-02609-x

23. Kavousi S, Hosseinpour A, Bahmanzadegan Jahromi F, Attar A. Efficacy of mesenchymal stem cell transplantation on major adverse cardiovascular events and cardiac function indices in patients with chronic heart failure: a meta-analysis of randomized controlled trials. J Transl Med. 2024;22:786. doi: 10.1186/s12967-024-05352-y

24. Kelly K, Bloor AJC, Griffin JE, Radia R, Yeung DT, Rasko JEJ. Two-year safety outcomes of iPS cell-derived mesenchymal stromal cells in acute steroid-resistant graft-versus-host disease. Nat Med. 2024;30:1556–1558. doi: 10.1038/s41591-024-02990-z

25. Sharma A, Gupta S, Archana S, Verma RS. Emerging Trends in Mesenchymal Stem Cells Applications for Cardiac Regenerative Therapy: Current Status and Advances. Stem Cell Rev Rep. 2022;18:1546–1602. doi: 10.1007/s12015-021-10314-8

26. Peng Y, Pan W, Ou Y, Xu W, Kaelber S, Borlongan CV, Sun M, Yu G. Extracardiac-Lodged Mesenchymal Stromal Cells Propel an Inflammatory Response Against Myocardial Infarction via Paracrine Effects. Cell Transplant. 2016;25:929–935. doi: 10.3727/096368915X689758

27. Deszcz I. Stem Cell-Based Therapy and Cell-Free Therapy as an Alternative Approach for Cardiac Regeneration. Stem Cells Int. 2023;2023:2729377. doi: 10.1155/2023/2729377

28. Hwang HJ, Chang W, Song BW, Song H, Cha MJ, Kim IK, Lim S, Choi EJ, Ham O, Lee SY, et al. Antiarrhythmic potential of mesenchymal stem cell is modulated by hypoxic environment. J Am Coll Cardiol. 2012;60:1698–1706. doi: 10.1016/j.jacc.2012.04.056

29. Yeghiazarians Y, Zhang Y, Prasad M, Shih H, Saini SA, Takagawa J, Sievers RE, Wong ML, Kapasi NK, Mirsky R, et al. Injection of bone marrow cell extract into infarcted hearts results in functional improvement comparable to intact cell therapy. Mol Ther. 2009;17:1250–1256. doi: 10.1038/mt.2009.85

30. Alrefai MT, Tarola CL, Raagas R, Ridwan K, Shalal M, Lomis N, Paul A, Alrefai MD, Prakash S, Schwertani A, et al. Functional Assessment of Pluripotent and Mesenchymal Stem Cell Derived Secretome in Heart Disease. Ann Stem Cell Res. 2019;2:29–36.

31. Angeli FS, Zhang Y, Sievers R, Jun K, Yim S, Boyle A, Yeghiazarians Y. Injection of human bone marrow and mononuclear cell extract into infarcted mouse hearts results in functional improvement. Open Cardiovasc Med J. 2012;6:38–43. doi: 10.2174/1874192401206010038

32. Karbassi E, Yoo D, Martinson AM, Yang X, Reinecke H, Regnier M, Murry CE. Noncontractile Stem Cell-Cardiomyocytes Preserve Post-Infarction Heart Function. Circ Res. 2024;135:967–969. doi: 10.1161/CIRCRESAHA.124.325133

33. Burt RK, Chen YH, Verda L, Lucena C, Navale S, Johnson J, Han X, Lomasney J, Baker JM, Ngai KL, et al. Mitotically inactivated embryonic stem cells can be used as an in vivo feeder layer to nurse damaged myocardium after acute myocardial infarction: a preclinical study. Circ Res. 2012;111:1286–1296. doi: 10.1161/CIRCRESAHA.111.262584

34. Thum T, Bauersachs J, Poole-Wilson PA, Volk HD, Anker SD. The dying stem cell hypothesis: immune modulation as a novel mechanism for progenitor cell therapy in cardiac muscle. J Am Coll Cardiol. 2005;46:1799–1802. doi: 10.1016/j.jacc.2005.07.053

35. Schmuck EG, Koch JM, Centanni JM, Hacker TA, Braun RK, Eldridge M, Hei DJ, Hematti P, Raval AN. Biodistribution and Clearance of Human Mesenchymal Stem Cells by Quantitative Three-Dimensional Cryo-Imaging After Intravenous Infusion in a Rat Lung Injury Model. Stem Cells Transl Med. 2016;5:1668–1675. doi: 10.5966/sctm.2015-0379

36. van den Akker F, Feyen DA, van den Hoogen P, van Laake LW, van Eeuwijk EC, Hoefer I, Pasterkamp G, Chamuleau SA, Grundeman PF, Doevendans PA, et al. Intramyocardial stem cell injection: go(ne) with the flow. Eur Heart J. 2017;38:184–186. doi: 10.1093/eurheartj/ehw056

37. Mitchell AJ, Sabondjian E, Sykes J, Deans L, Zhu W, Lu X, Feng Q, Prato FS, Wisenberg G. Comparison of initial cell retention and clearance kinetics after subendocardial or subepicardial injections of endothelial progenitor cells in a canine myocardial infarction model. J Nucl Med. 2010;51:413–417. doi: 10.2967/jnumed.109.069732

38. Hodgson-Garms M, Moore MJ, Martino MM, Kelly K, Frith JE. Proteomic profiling of iPSC and tissue-derived MSC secretomes reveal a global signature of inflammatory licensing. NPJ Regen Med. 2025;10:7. doi: 10.1038/s41536-024-00382-y

39. Romanazzo S, Kopecky C, Jiang S, Doshi R, Mukund V, Srivastava P, Rnjak-Kovacina J, Kelly K, Kilian KA. Biomaterials directed activation of a cryostable therapeutic secretome in induced pluripotent stem cell derived mesenchymal stromal cells. J Tissue Eng Regen Med. 2022;16:1008–1018. doi: 10.1002/term.3347

40. Zhang Y, Zhang Z, Gao F, Tse HF, Tergaonkar V, Lian Q. Paracrine regulation in mesenchymal stem cells: the role of Rap1. Cell Death Dis. 2015;6:e1932. doi: 10.1038/cddis.2015.285

41. Valencia J, Yanez RM, Muntion S, Fernandez-Garcia M, Martin-Rufino JD, Zapata AG, Bueren JA, Vicente A, Sanchez-Guijo F. Improving the therapeutic profile of MSCs: Cytokine priming reduces donor-dependent heterogeneity and enhances their immunomodulatory capacity. Front Immunol. 2025;16:1473788. doi: 10.3389/fimmu.2025.1473788

