## Supplementary Materials for "Device encapsulated MSCs for adaptive secretome therapy to effectively target ischaemic heart injury"

### Methods

**Cell culture**

Cardiomyocyte differentiation from iPSCs: Cardiomyocytes were derived from human iPSCs as previously described with modifications ^1^. Briefly, iPSCs were seeded onto Matrigel (Corning, NY, USA) coated plates at a density of 1.25×10^5^ cells/cm^2^ in TeSR-E8 medium supplemented with 10 μM Y-27632 (Abcam, Cambridge, UK). After 48 hours when the cells were 100% confluent, which is referred to as day 0, medium was replaced with RPMI 1640 basal medium (Thermo Fisher Scientific) containing B-27 without insulin supplement (Thermo Fisher Scientific), growth factor reduced Matrigel (1:60 dilution) and 10 μM CHIR99021 (STEMCELL Technologies). On day 1, medium was replaced with RPMI 1640 basal medium containing B-27 without insulin supplement. On day 2, medium was changed to RPMI 1640 basal medium containing B-27 without insulin supplement and 5 μM IWP2 (Sigma-Aldrich) for 72 hours. From day 5 onwards, cells were cultured in RPMI 1640 basal medium containing B-27 supplement (Thermo Fisher Scientific) and 200 μg/mL L-ascorbic acid 2-phosphate sesquimagnesium salt hydrate (Sigma-Aldrich), referred to as cardiomyocyte medium, and medium was changed every 2-3 days. On day 12, cardiomyocytes were dissociated into single cells and split at 1:4 ratio onto Matrigel coated plates in cardiomyocyte replating medium (DMEM/F-12 GlutaMAX medium (Thermo Fisher Scientific) supplemented with 20% fetal bovine serum (Bovogen Biologicals, Victoria, Australia), 0.1 mM 2-mercaptoethanol, 0.1 mM nonessential amino acids, 50 U/mL antibiotic-antimycotic (Thermo Fisher Scientific) and 10 μM Y-27632). On day 13, medium was changed to cardiomyocyte medium. From days 14–19, cardiomyocytes were enriched to >90% cardiac troponin T positive cells by culture in glucose-free DMEM medium (Thermo Fisher Scientific) containing 4 mM lactate (Sigma-Aldrich).

Endothelial cell differentiation from iPSCs: Human iPSCs were differentiated into CD31 positive endothelial cells according to a previously published method ^2^. For endothelial differentiation, iPSCs were dissociated into single cells and seeded onto Matrigel-coated plates at a density of 1×10^5^ cells/cm^2^ in TeSR-E8 medium supplemented with 10 μM Y-27632. After 24 hours, referred to as day 0, medium was replaced with DMEM/F12 GlutaMAX medium containing N-2 supplement (Thermo Fisher Scientific), B-27 supplement, 8 µM CHIR99021 and 25 ng/mL BMP4 (STEMCELL Technologies) for 3 days. Medium was then replaced with StemPro-34 SFM complete medium (Thermo Fisher Scientific) supplemented with 200 ng/mL VEGF-165 (PeproTech, NJ, USA) and 2 µM forskolin (Sigma-Aldrich) for 3 days. On day 6, CD31 positive cells were sorted by FACS using a CD31-conjugated antibody (BD Pharmingen, NJ, USA) and expanded on human plasma fibronectin (Thermo Fisher Scientific) coated plates and cultured in EGM2-MV medium (Lonza, Basel, Switzerland) supplemented with 50 ng/mL VEGF-165.

Smooth muscle cell differentiation from iPSCs: Contractile vascular smooth muscle cells were differentiated from human iPSCs according to a published protocol with modifications ^3^. iPSCs were dissociated and seeded onto Matrigel-coated plates at a density of 4×10^5^ cells/cm^2^ in TeSR-E8 medium supplemented with 10 μM Y-27632. After 24 hours, referred to as day 0, medium was replaced with N2B27 medium (1:1 ratio mixes of DMEM/F-12 GlutaMAX medium and Neurobasal medium (Thermo Fisher Scientific) plus N-2 supplement and B-27 minus vitamin A supplement (Thermo Fisher Scientific)) supplemented with 8 μM CHIR99021 and 25 ng/mL BMP4. On day 3, medium was replaced with N2B27 medium supplemented with 10 ng/mL PDGF-BB (PeproTech) and 2 ng/mL Activin A (PeproTech). On day 4, cells were replated at 8×10^5^ cells/cm^2^ on collagen (Sigma-Aldrich) coated plates in N2B27 medium supplemented with 2 μg/mL heparin (Sigma-Aldrich) and 2 ng/mL Activin A until day 7 and then replaced with SmGM-2 medium (Lonza) until day 10.

Cardiac fibroblast differentiation from iPSCs: Cardiac fibroblasts were differentiated according to a published method ^4^. iPSCs were dissociated into single cells and seeded onto Matrigel-coated plates at a density of 2.5×10^4^ cells/cm^2^ in TeSR-E8 medium supplemented with 10 μM Y-27632. After 24 hours, referred to as day 0, medium was replaced with STEMdiff™ APEL™ medium (STEMCELL Technologies) containing 20 ng/ml BMP4, 20 ng/ml Activin A and 1.5 µM CHIR99021 for 3 days. On day 3, medium was replaced with STEMdiff™ APEL™ medium containing 30 ng/ml BMP4, 1 µM retinoic acid (Sigma-Aldrich) and 5 µM IWP-2. On day 6, medium was replaced with STEMdiff™ APEL™ medium containing 30 ng/ml BMP4 and 1 µM retinoic acid (Sigma-Aldrich). On day 9, cells were dissociated and replated at 1.5×10^4^ cells/cm^2^ on human fibronectin-coated plates and cultured in STEMdiff™ APEL™ medium containing 10 μM SB431542 (Stem Cell Technologies) and 10 μM Y-27632. On day 13, epicardial cells were dissociated and replated at 2.5×10^4^ cells/cm^2^ on 0.1% porcine gelatine-coated plates and cultured in STEMdiff™ APEL™ medium containing 10 ng/ml bFGF (PeproTech) and 10 μM Y-27632, and medium was changed every 2 days without Y-27632. From day 20 to 30, the medium was replaced with changed to FGM-3 medium (Lonza) and replaced every 2-3 days.

**Engineered** **cardiac microtissue**

To construct the multicellular cardiac microtissue, iPSC-derived cardiomyocytes (3.5×10^4^ cells), endothelial cells (1.25x10^4^ cells), smooth muscle cells (1.25 x10^3^ cells) and cardiac fibroblasts (1.25x10^3^ cells) were seeded per well into ultra-low attachment round-bottom 96-well plates in spheroid induction medium (cardiomyocyte replating medium, EGM2-MV, SmGM-2, and FGM-3 at a 1:1:1:1 ratio and supplemented with 50 ng/mL VEGF-165 and 10 μM Y-27632), and spun at 150 *g* for 3 minutes. After 48 hours, medium was changed to cardiac spheroid maintaining medium (cardiomyocyte medium, EGM2-MV, SmGM-2, and FGM-3 at a 1:1:1:1 ratio and supplemented with 50 ng/mL VEGF-165). After a further 24 hours, compacted spheroids were embedded in 10 µL growth factor reduced Matrigel and transferred to tissue culture plates coated with anti-adherence rinsing solution (STEMCELL Technologies) containing cardiac spheroid maintaining medium. Engineered cardiac microtissues were maintained at 37°C in a humidified 5% CO_2_ incubator on an orbital shaker rotating at 60 rpm.

**Cancer cells**

MDA-MB-231 breast cancer cells were cultured in DMEM medium (high glucose; Sigma-Aldrich) supplemented with 2 mM GlutaMAX (Thermo Fisher Scientific) and 10% foetal bovine serum (Sigma-Aldrich). MG63 osteosarcoma cells were cultured in alpha-MEM medium (Thermo Fisher Scientific) supplemented with 2 mM GlutaMAX and 10% foetal bovine serum. Cells were maintained at 37°C in a humidified 5% CO_2_ incubator and media was changed every 2-3 days. For treatment, cells were seeded into 96-well black optiplate (Corning, NY, USA) at a density of 9.38x10^3^/cm^2^ in their respective culture media. After 24 hours, cells were treated with conditioned media collected from Cymerus MSCs for 48 hours. Cell viability was assessed using the CellTiter-Blue assay (Promega, WI, USA) according to the manufacturer’s protocol. The CellTiter-Blue assay quantifies viable cells based on the conversion of resazurin to resorufin, a fluorescent product generated by metabolically active cells. Fluorescent intensity was measured at 560/590 nm (excitation and emission) using the EnSpire Multimode Plate Reader (PerkinElmer, MA, USA). All readings were subtracted for the background fluorescence (medium with CellTiter-Blue reagent without cells), and cell viability was expressed as fold change relative to control using the formula: Cell viability (fold change to control) = sample fluorescence / control fluorescence.

**Echocardiography**

Transthoracic echocardiography was performed at baseline (before surgery) and on day 1, 7, 28, 56 and 84 post-surgery under light anaesthesia with 2% isoflurane, using either a Vivid 7 Dimension ultrasound imaging machine with a 10 MHz phased array probe or a Vivid E9 ultrasound imaging machine with a 12 MHz phased array probe (GE HealthCare, IL, USA) ^5^. Two-dimensional parasternal long-axis views of the left ventricle were obtained for measurement of ejection fraction (EF) offline using EchoPAC software (GE HealthCare) for left ventricular dimension and ejection fraction. All parameters were assessed using an average of three consecutive beats and calculations were made in accordance with the American Society of Echocardiography guidelines ^6^.

**Blood biochemistry**

Blood samples were collected on days 0, 7, 28, 56 and 84 via tail prick to measure blood glucose concentrations using a handheld glucometer (Accu-Chek Performa, Roche, Basel, Switzerland). In Part 2 validation study, an additional blood sample for blood glucose measurement was collected before initiating the high fat diet. At tissue harvest on day 84 post-surgery, blood was collected from the abdominal aorta following echocardiography in EDTA tubes, kept on ice, and centrifuged at 3000 rpm at 4°C. Plasma aliquots were frozen for subsequent biochemical analysis of lipids and cytokines. A separate EDTA whole blood sample was also aliquoted and frozen for HbA1c analysis. Frozen samples were thawed to measure total cholesterol, LDL-cholesterol, HDL-cholesterol, triglycerides and HbA1c using a Cobas integra 400 Plus® chemical analyser (Roche Diagnostics, NSW, Australia) with manufacturer-provided standards and controls ^7^. An aliquot of plasma was also thawed for the analysis of inflammatory markers and cytokines C-reactive protein, Interleukin-1β (IL-1β), IL-6 and tumour necrosis factor-α (TNFα) were analysed using a ProcartaPlex Mix and Match Rat panel (Invitrogen, MA, US) and measured on a Bio-Plex 200 system (Bio-Rad, NSW, Australia). Samples were measured in duplicate in accordance with the manufacturer’s description, and the analysed data was obtained from the Bio-Plex Manager Software (Bio-Rad).

**Histology**

At 84 days post-surgery, the hearts and devices of anesthetized animals (2-3% isoflurane in oxygen) were excised, weighed and sliced into 5 equal short-axis transverse ventricular slices from the apex to the base. Heart sections were then fixed in 10% neutral buffer formalin and processed for paraffin embedding and histochemical stains. To determine the infarct size, picrosirius red stained histology images of all five ventricular slices were captured. Infarct size was calculated as the percentage of the area occupied by scar divided by the total left ventricular area. To determine the myocardial interstitial fibrosis, 10 random histological fields of picrosirius red stained sections (remote region) were captured at 400 times magnification. Mean cardiomyocyte cross-sectional area was determined in the non-infarcted remote region stained with wheat germ agglutinin conjugated with rhodamine. Wheat germ agglutinin-stained cell membrane of 250 cardiomyocytes (positioned perpendicular to the plane of the section with cell membrane clearly outlined, endomyocardium region) from 5 random endocardial fields at 200 times magnification were sampled from each heart using the BX-61 Olympus fluorescence microscope. The total vascular density including the capillaries was analysed on tissue sections stained with *Griffonia simplicifolia* lectin I. Five high-power fields in the non-infarcted remote left ventricular (endomyocardium region) from each heart were randomly captured at 400 times magnification.

Devices were fixed in 10% neutral buffer formalin and processed for paraffin embedding, and 5 µm tissue sections were mounted on Trajan Microscope T7611 Series 3 Adhesive slides (Trajan Scientific Australia Pty Ltd). Tissue morphology was evaluated using haematoxylin and eosin staining, while Alcian Blue staining was used to identify cartilaginous tissue. Vascularization of the transplanted device was assessed by staining with *Griffonia simplicifolia* lectin I. Cell fate was analysed through immunostaining with antibodies against CD31 (4 μg/mL, mouse monoclonal, clone JC70A, Dako), cleaved caspase-3 (0.27 µg/mL, rabbit monoclonal, clone 5A1E, Cell Signaling Technology), Ki67 (0.3 μg/mL, rabbit monoclonal, clone SP6, Abcam), and the human-specific KU80 (0.12 μg/mL, rabbit monoclonal, clone EPR3468, Abcam).

**Collection of conditioned medium**

Cymerus MSCs were cultured to 90% confluence in T150 cm^2^ culture flasks and cultured in 30 mL of Cymerus MSC serum-free expansion medium for 3 days at 37ºC prior to harvesting the conditioned medium. The conditioned medium was centrifuged at 2000 *g* for 10 minutes at 4ºC and the supernatant was collected and concentrated using the Amicon Ultra-15 centrifugal filter devices with 3000 molecular weight cut off membrane (Millipore, MA, USA). The protein concentration of concentrated conditioned medium was determined using the Pierce BCA protein assay kit (Thermo Fisher Scientific).

**Simulated IRI (*in vitro*)**

Engineered cardiac microtissues were subjected to 3 hours of hypoxia and 24 hours of reoxygenation to simulate IRI. Hypoxia was induced in a hypoxic chamber (STEMCELL Technologies) where oxygen was purged by pure nitrogen gas for 15 minutes and using a buffer simulating the conditions of ischaemia (in mmol/L: 1.0 KH_2_PO_4_, 10.0 NaHCO_3_, 1.2 MgCl_2_.6H_2_0, 25.0 Na(4-(2-hydroxyethyl)-1-piperazineethanesulfonic acid) (HEPES), 74.0 NaCl, 16.0 KCl, 1.2 CaCl_2_ and 10 mM 2-deoxyglucose, pH 6.7), gassed with pure nitrogen gas for 5 minutes. Reoxygenation was achieved by replacing the buffer with spheroid medium and culturing in a humidified incubator at 37ºC (~21% O_2_). Engineered cardiac microtissues cultured in spheroid medium at 37ºC in a humidified CO_2_ incubator throughout the hypoxia and reoxygenation period served as the normoxic control group. Engineered cardiac microtissues were randomly assigned to receive concentrated Cymerus MSC serum-free expansion medium (as vehicle control) or concentrated conditioned media of Cymerus MSCs at reoxygenation for 24 hours.

**Contractility of engineered cardiac microtissue**

At the end of the 24-hour reoxygenation period, brightfield videos of contracting engineered cardiac microtissues were captured at 40x magnification and 60 frames/second using an IX-71 microscope (Olympus) coupled with a DP74 camera (Olympus). Videos were analyzed using the MUSCLEMOTION software tool ^8^ in ImageJ using the following settings: frames/second (60), speedWindow (2), default noise reduction, automatic reference frame and automatic peak fitting. Normalized beat rate variability was calculated as the Root Mean Square of Successive Differences (RMSSD) divided by the R-R interval, where RMSSD=$\sqrt{\frac{\sum_{i=1}^{N-1} {({RR}_{i}-{RR}_{i+1})}^{2}}{N-1}}$, *N* is the number of total beats, and *RR* represents the time difference between adjacent peaks ^1^.

**Viability of engineered cardiac microtissue**

The CellTiter-Glo luminescent cell viability assay was performed according to the manufacturer’s protocol (Promega) with modifications. Briefly, the media was removed from each well of the 96-well plate. Then, 100 μL of mixed dye (50 μL of cell culture media + 50 μL of dye) was added to each well. The plate was then shaken vigorously for 5 minutes to induce cell lysis, followed by a 20-minute incubation at room temperature to stabilise the luminescent signal. Luminescence was recorded using the POLARstar plate reader (BMG Labtech, VIC, Australia). All readings were corrected for the background signals (CellTiter-Glo reagent + cell culture media with no cells) and cell viability was expressed as a fold change relative to the normoxia control group.

**Mitochondrial reactive oxygen species (ROS)**

At the end of the 24 hours reoxygenation, mitochondrial ROS levels were assessed using the MitoSOX™ Red dye (Thermo Fisher Scientific). Engineered cardiac microtissues were stained with 5 μM MitoSOX™ Red for 30 minutes in HBSS++ at 37°C in a humidified CO_2_ incubator, washed once with HBSS++ and then imaged immediately in HBSS++ solution. Fluorescence was acquired at 50x magnification to capture the entire engineered cardiac microtissue on a Thunder microscope (Leica). The fluorescence intensity of each engineered cardiac microtissue was quantified in ImageJ to determine the corrected total cell fluorescence ^9^.

**Immunofluorescent staining of engineered cardiac microtissue**

Engineered cardiac microtissues were fixed in 10% neutral buffered formalin for 1 hour at room temperature and then dehydrated in 20% sucrose solution for 24 hours. Dehydrated samples were embedded in Optimal Cutting Temperature compound (Sakura Finetek, Tokyo, Japan) and cryosections (6 µm thick) were treated with 0.2% Triton X-100 permeabilization buffer and Protein Block (Dako, Victoria, Australia) before staining with primary antibodies; cardiac troponin T (2 μg/mL, rabbit polyclonal, Abcam), cardiac troponin T (4 μg/mL, mouse monoclonal, clone 1C11, Abcam), CD31 (4 μg/mL, mouse monoclonal, clone JC70A, Dako), SM22 (5 μg/mL, rabbit polyclonal, Abcam), or vimentin (0.3 μg/mL, mouse monoclonal, clone V9, Dako), followed by Alexa Fluor-488-conjugated goat-anti-rabbit or goat-anti-mouse (10 μg/mL, Invitrogen) and Alexa Fluor-594-conjugated goat-anti-mouse or goat-anti-rabbit (10 μg/mL, Invitrogen) secondary antibodies. Sections were then counterstained with 1 μg/mL of DAPI (Invitrogen) for nuclear staining. Epifluorescence images of immunostained sections were acquired with an Olympus BX61 upright microscope using analySIS software.

**Mass spectrometry-based proteomics**

For proteomic analysis of encapsulated Cymerus MSCs (cell lysate) and their secretome (conditioned media, CM), samples were solubilized in 1% (v/v) sodium dodecyl sulphate (SDS) containing 50 mM HEPES (pH 8.0) and HALT protease and phosphatase inhibitor (#78442, Thermo Fisher Scientific). Further, global cell proteomics was performed on a monolayer of Cymerus MSCs (2D) and human iPSCs (iPS-Foreskin-2 cell line) for comparative purpose. Protein lysates were further homogenized by tip-probe sonication on ice and quantified by microBCA (Thermo Fisher Scientific) ^10^.

Cell and secretome samples (10 µg protein) were reduced and alkylated with 10 mM DTT and 20 mM IAA as described^37^. Briefly, samples were prepared using SP3 protocol ^11^ and digested using Trypsin and Lys-C (1:50 and 1:100 enzyme-to-protein ratio, respectively) overnight at 37°C. Samples were acidified after digestion to final concentration of 2% formic acid before vacuum lyophilisation. Samples were reconstituted in 10 µL of 0.07% (v/v) trifluoroacetic acid in LC-MS grade water, peptides quantified using fluorometric peptide assay (Thermo Fisher Scientific). LC-MS data acquisition was performed on Q Exactive HF-X benchtop Orbitrap mass spectrometer coupled with UltiMate™ NCS-3500RS nano-HPLC and operated with Xcalibur software as previously described ^10,12^.

For proteomics analyses, we investigated biological replicates for global cell and secretome proteomics: encapsulated Cymerus MSCs pre-implantation (n=2), encapsulated Cymerus MSCs post-implantation (*ex vivo* cultured) (n=3), 2D Cymerus MSCs monolayer (n=2), iPSCs (n=3). MS-based proteomics data is deposited to the ProteomeXchange Consortium via the MassIVE partner repository and available via MassIVE with identifier (MSV000097257).

DIA-MS spectra were processed using DIA-NN software (v1.9) ^13^ as previously reported ^10,14^. The DIA-MS spectra were searched against human proteome database (UP000005640, #83,401). Perseus (v2.0.11) was applied for data processing and analysis, with scatter plots/bar charts generated using GraphPad Prism (v8.0.1) or Microsoft Excel. Protein intensities were log2 transformed and normalized using quantile normalization. Proteins were subjected to PCA and unpaired student’s t-test or Welch’s T-test with missing values imputed from normal distribution (width 0.3, downshift 1.8). g:Profiler and Reactome pathway databases were utilized for Gene Ontology functional enrichment and network/pathway analysis, significance P < 0.05.

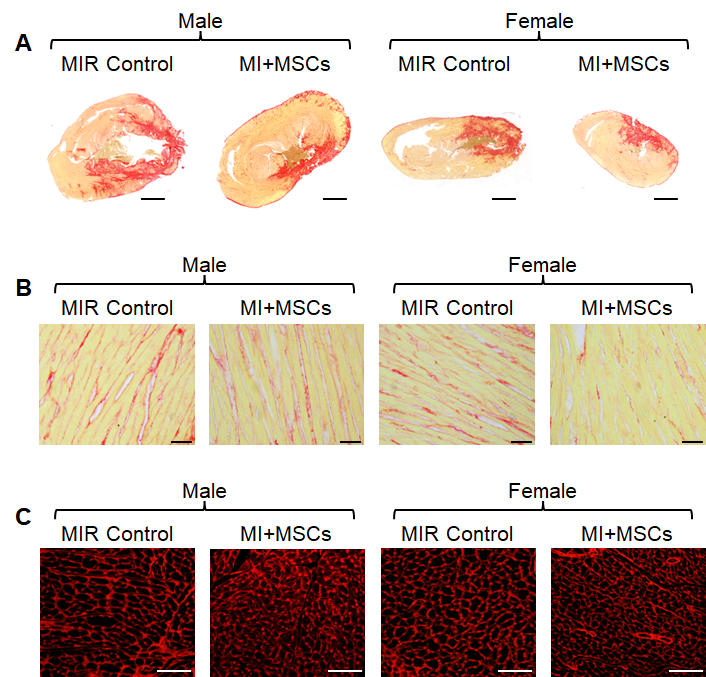

**Supplementary Figure S1**. **Histomorphological changes of aged rat heart tissues.** Representative cross-sectional images of myocardium collected 84 days post-surgery from rats (aged 51-53 weeks) subjected to sham operation, reperfused myocardial infarction (MIR control) and reperfused myocardial infarction treated with 4x10^6^ Cymerus MSCs encapsulated in a Procyon device. Tissue sections were stained with picrosirius red (A-B) or wheat germ agglutinin (C). Scale bar = 2mm (A), 50µm (B), and 100µm (C).

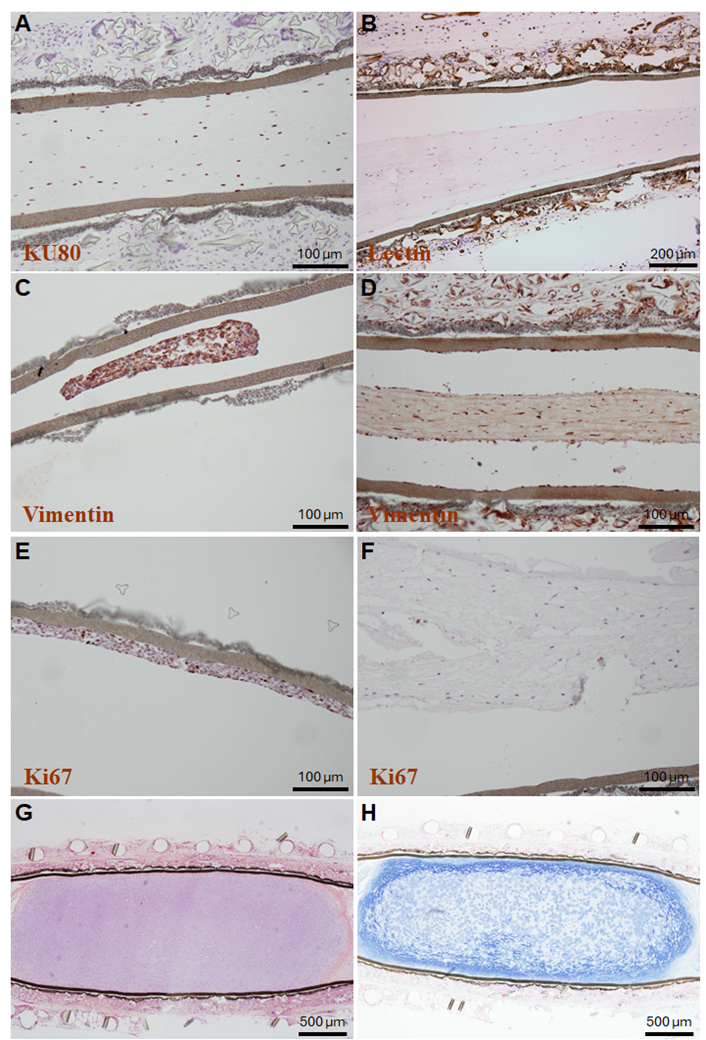

**Supplementary Figure S2**. **Morphological changes of Cymerus MSCs encapsulated within a Procyon immunoisolation device.** Representative images of encapsulated Cymerus MSCs stained with human-specific KU80 (A), rodent-specific lectin (B), vimentin (C-D), Ki67 (E-F), haematoxylin and eosin (G), and Alcian blue (H), shown before (C, E) and after (A, B, D, F) *in vivo* implantation in middle-aged male and female rats following reperfused myocardial infarction.

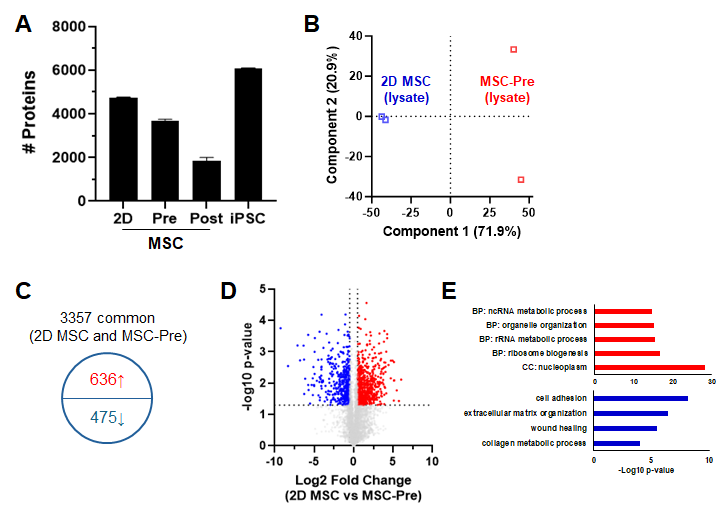

**Supplementary Figure S3**. (**A**) Quantitative proteomic profiling reveals in-depth coverage and identified protein in 2D Cymerus MSCs (2D MSC), encapsulated Cymerus MSCs at 0 (MSC-Pre) and 12 (MSC-Post) weeks post-implantation, and human iPSCs. (**B**) Principal component analysis of indicated cellular proteomes of 2D Cymerus MSC and MSC-Pre. **(C)** Venn diagram of the distribution of identified proteins, their similarity and differential expression. (**D)** Volcano plot of the differentially expressed proteins in cellular proteome of 2D MSC relative to MSC-Pre. **(E)** Enrichment analysis of GO networks/terms (P < 0.05) in proteins significantly enriched in cellular proteome of 2D MSC (red) and in cellular proteome of MSC-Pre (blue).

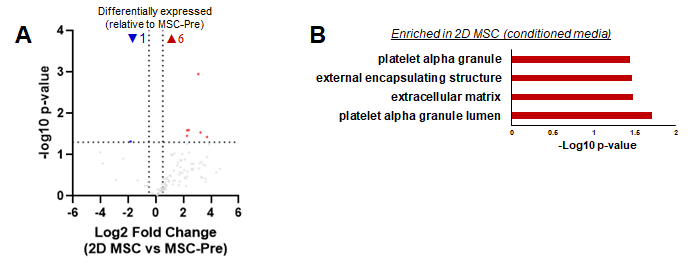

**Supplementary Figure S4.** (**A)** Volcano plot of the differentially expressed proteins in conditioned media (secretome) collected from 2D Cymerus MSC (2D MSC) relative to encapsulated Cymerus MSCs at pre-implantation (MSC-Pre). **(B)** Enrichment analysis of GO networks/terms (P < 0.05) in proteins significantly enriched in conditioned media of 2D MSCs (red).

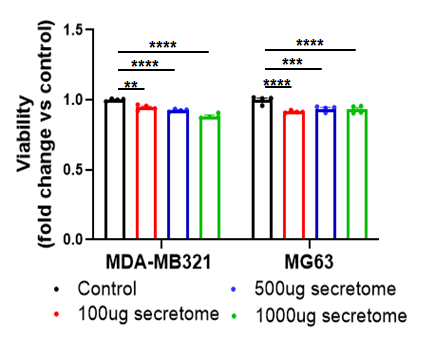

**Supplementary Figure S5. Effect of Cymerus-MSC secretome on human cancer cell growth.** Cymerus-MSC secretome significantly reduced viability of human breast cancer (MDA-MB321) and osteosarcoma (MG63) cells. n = 4 replicates. Data are presented as mean ± SEM. **P < 0.01, ***P < 0.001, ****P < 0.0001 by one-way ANOVA with Bonferroni post hoc test.

1.	Lyu Q, Gong S, Lees JG, Yin J, Yap LW, Kong AM, Shi Q, Fu R, Zhu Q, Dyer A, et al. A soft and ultrasensitive force sensing diaphragm for probing cardiac organoids instantaneously and wirelessly. *Nat Commun*. 2022;13:7259. doi: 10.1038/s41467-022-34860-y

2.	Kong AM, Yap KK, Lim SY, Marre D, Pebay A, Gerrand YW, Lees JG, Palmer JA, Morrison WA, Mitchell GM. Bio-engineering a tissue flap utilizing a porous scaffold incorporating a human induced pluripotent stem cell-derived endothelial cell capillary network connected to a vascular pedicle. *Acta Biomater*. 2019;94:281-294. doi: 10.1016/j.actbio.2019.05.067

3.	Patsch C, Challet-Meylan L, Thoma EC, Urich E, Heckel T, O'Sullivan JF, Grainger SJ, Kapp FG, Sun L, Christensen K, et al. Generation of vascular endothelial and smooth muscle cells from human pluripotent stem cells. *Nat Cell Biol*. 2015;17:994-1003. doi: 10.1038/ncb3205

4.	Giacomelli E, Meraviglia V, Campostrini G, Cochrane A, Cao X, van Helden RWJ, Krotenberg Garcia A, Mircea M, Kostidis S, Davis RP, et al. Human-iPSC-Derived Cardiac Stromal Cells Enhance Maturation in 3D Cardiac Microtissues and Reveal Non-cardiomyocyte Contributions to Heart Disease. *Cell Stem Cell*. 2020;26:862-879 e811. doi: 10.1016/j.stem.2020.05.004

5.	Kompa AR, Greening DW, Kong AM, McMillan PJ, Fang H, Saxena R, Wong RCB, Lees JG, Sivakumaran P, Newcomb AE, et al. Sustained subcutaneous delivery of secretome of human cardiac stem cells promotes cardiac repair following myocardial infarction. *Cardiovasc Res*. 2021;117:918-929. doi: 10.1093/cvr/cvaa088

6.	Schiller NB, Shah PM, Crawford M, DeMaria A, Devereux R, Feigenbaum H, Gutgesell H, Reichek N, Sahn D, Schnittger I, et al. Recommendations for quantitation of the left ventricle by two-dimensional echocardiography. American Society of Echocardiography Committee on Standards, Subcommittee on Quantitation of Two-Dimensional Echocardiograms. *J Am Soc Echocardiogr*. 1989;2:358-367. doi: 10.1016/s0894-7317(89)80014-8

7.	Savira F, Wang BH, Edgley AJ, Jucker BM, Willette RN, Krum H, Kelly DJ, Kompa AR. Inhibition of apoptosis signal-regulating kinase 1 ameliorates left ventricular dysfunction by reducing hypertrophy and fibrosis in a rat model of cardiorenal syndrome. *Int J Cardiol*. 2020;310:128-136. doi: 10.1016/j.ijcard.2020.04.015

8.	Sala L, van Meer BJ, Tertoolen LGJ, Bakkers J, Bellin M, Davis RP, Denning C, Dieben MAE, Eschenhagen T, Giacomelli E, et al. MUSCLEMOTION: A Versatile Open Software Tool to Quantify Cardiomyocyte and Cardiac Muscle Contraction In Vitro and In Vivo. *Circ Res*. 2018;122:e5-e16. doi: 10.1161/CIRCRESAHA.117.312067

9.	Lees JG, Gardner DK, Harvey AJ. Nicotinamide adenine dinucleotide induces a bivalent metabolism and maintains pluripotency in human embryonic stem cells. *Stem Cells*. 2020;38:624-638. doi: 10.1002/stem.3152

10.	Cross J, Rai A, Fang H, Claridge B, Greening DW. Rapid and in-depth proteomic profiling of small extracellular vesicles for ultralow samples. *Proteomics*. 2024;24:e2300211. doi: 10.1002/pmic.202300211

11.	Hughes CS, Moggridge S, Muller T, Sorensen PH, Morin GB, Krijgsveld J. Single-pot, solid-phase-enhanced sample preparation for proteomics experiments. *Nat Protoc*. 2019;14:68-85. doi: 10.1038/s41596-018-0082-x

12.	Tham YK, Bernardo BC, Claridge B, Yildiz GS, Woon LM, Bond S, Fang H, Ooi JYY, Matsumoto A, Luo J, et al. Estrogen receptor alpha deficiency in cardiomyocytes reprograms the heart-derived extracellular vesicle proteome and induces obesity in female mice. *Nat Cardiovasc Res*. 2023;2:268-289. doi: 10.1038/s44161-023-00223-z

13.	Demichev V, Messner CB, Vernardis SI, Lilley KS, Ralser M. DIA-NN: neural networks and interference correction enable deep proteome coverage in high throughput. *Nat Methods*. 2020;17:41-44. doi: 10.1038/s41592-019-0638-x

14.	Fang H, Greening DW. An Optimized Data-Independent Acquisition Strategy for Comprehensive Analysis of Human Plasma Proteome. *Methods Mol Biol*. 2023;2628:93-107. doi: 10.1007/978-1-0716-2978-9_7
